# Human ribosomes interactions reveal the binding landscape of neomycin

**DOI:** 10.1101/2025.07.22.666027

**Authors:** Meng Wu, Lingfan Zhu, Keqin Wu, Changfeng Jia, Xinyi Zhang, Zhongcheng Zhao, Haihan Liu, Chuansen Yi, Shuangli Sun, Ke Liu, Zhe Zhang, Lei Guo, Xuben Hou, Wenfei Li

## Abstract

Aminoglycosides such as neomycin are clinically limited by toxicity, commonly attributed to the inhibition of mitoribosomes. Here, we investigate this model by integrating Cryo-EM and functional assays to assess neomycin’s binding landscape and functional consequences for human ribosomes. Cryo-EM reveals extensive multi-site binding of neomycin on both human mitochondrial and cytosolic ribosomes *in vitro*, including conserved regions of the decoding center. In contrast, ribosomes isolated from neomycin-treated cells show markedly different patterns, with no detectable neomycin density on the mitoribosomes and reduced occupancy at a subset of cytosolic ribosome sites. Functional assays indicate that neomycin remained active in cells, inducing widespread stop-codon readthrough while having minimal effects on global mitochondrial and cytosolic protein synthesis. Together, these findings highlight a disconnect between the structural binding landscape and ribosome-associated neomycin recovered following cellular exposure, while linking neomycin exposure to altered translational fidelity with minimal effects on global protein synthesis.

## INTRODUCTION

Precise control of protein synthesis is fundamental to cellular life, and its disruption represents a major mechanism of antibiotic action(Kohanski et al. 2010, Wilson 2014, Lin et al. 2018, Paternoga et al. 2023). Aminoglycosides are highly effective against bacteria, yet their clinical use is limited by dose-dependent ototoxicity and nephrotoxicity(Zaher and Green 2009, Becker and Cooper 2013). While bacterial selectivity is largely determined by structural features within the decoding center(Moazed and Noller 1987, Recht et al. 1999), the evolutionary conservation of ribosomal functional elements has raised long-standing concerns regarding off-target interactions with human ribosomes. In particular, the structural similarity between bacterial and mitochondrial ribosomes has led to the prevailing view that mitoribosome inhibition underlies aminoglycoside toxicity(Prezant et al. 1993, Böttger et al. 2001, Hobbie et al. 2008, Guan 2011, Matt et al. 2012, Amunts et al. 2015, Greber et al. 2015). In parallel, aminoglycosides have also been shown to affect eukaryotic cytosolic ribosomes, altering translational fidelity and promoting premature termination codon readthrough(Palmer et al. 1979, Manuvakhova et al. 2000, Keeling et al. 2014, Fosso et al. 2014, Prokhorova et al. 2017), a phenomenon recently visualized at high resolution in fungal 80S ribosomes(Kolosova et al. 2025) and explored therapeutically for nonsense mutation suppression(Shalev et al. 2015, Sabbavarapu et al. 2016).

Neomycin is a representative 4,5-linked aminoglycoside(Benveniste and Davies 1973, Fosso et al. 2014) that inhibits bacterial translation primarily through interactions with the decoding center(Wallis et al. 1995, Wang et al. 2012). In bacteria, binding to h44 of the small subunit (SSU) and H69 of the large subunit (LSU) promotes codon misreading and disrupts translational fidelity(Fourmy et al. 1996, Borovinskaya et al. 2007, Wang et al. 2012, Wasserman et al. 2015, Paternoga et al. 2023). Recent structural and biochemical studies have established that aminoglycoside activity depends on specific conformational rearrangements within both the drug and the ribosome, highlighting the dynamic nature of ribosome–drug recognition(Dey et al. 2025). This complexity is further highlighted by specialized translation factors that selectively rescue drug-corrupted ribosomes(Ghosh Dastidar et al. 2025). Together, these observations suggest that structural compatibility alone may be insufficient to predict physiological ribosome engagement, raising the possibility that interactions observed under *in vitro* conditions may not fully reflect those occurring within the cellular environment.

Despite extensive knowledge of aminoglycoside action in bacterial systems, how neomycin engages human ribosomes remains poorly understood. To address this question, we systematically investigated neomycin interactions with both human mitochondrial (55S) and cytosolic (80S) ribosomes. We combined high-resolution structural analysis and functional characterization to define the binding landscape observed *in vitro*, examine ribosome-associated drug binding recovered from neomycin-treated cells, and assess the functional consequences of neomycin treatment in human cells. This integrated approach provides a framework for evaluating aminoglycoside action in human cells.

## RESULTS

### Neomycin exhibits extensive and conformation-dependent binding on the human mitoribosome

To investigate the molecular basis of neomycin’s interaction with mitoribosomes, we performed cryo-EM on mitoribosomes rapidly isolated from HEK293F cells and incubated with 50 μM neomycin, a 4,5-linked aminoglycoside containing the conserved 2-deoxystreptamine (2-DOS) core (**Fig. 1A**). This yielded six maps with average resolutions ranging from 2.9−3.3 Å, representing distinct rotational and tRNA-occupancy conformations (**Supplementary Fig. S1, Supplementary Table S1**). These maps were of sufficient quality to resolve endogenous cofactors, including polyamines and Fe-S clusters, supporting confident assignment of neomycin-binding densities (**Supplementary Fig. S1C, D**). Across the six states, we identified nine distinct neomycin-binding sites (mtSSU Neo1-3, mtLSU Neo4-9), revealing an extensive interaction landscape across the mitoribosome (**Fig. 1B**, **Supplementary Table S2**).

**Figure 1.**
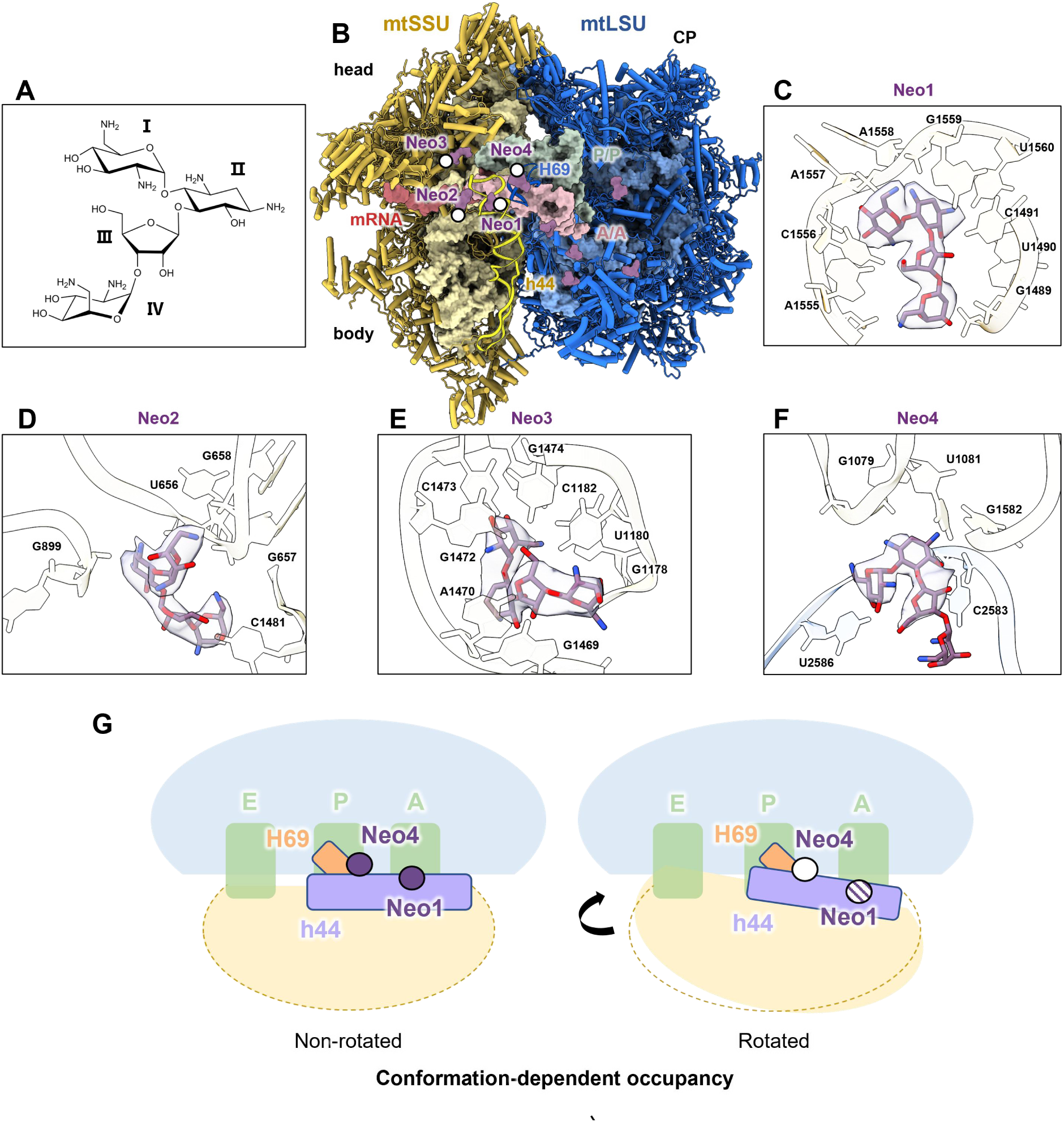
Structural analysis of neomycin binding to mitoribosome *in vitro*. **(A)** Chemical structure of neomycin. **(B)** Overall structure of the neomycin-bound mitoribosome in non-rotated state comprising the mtSSU (yellow), mtLSU (blue), mRNA (red), A-site tRNA (magenta) and P-site tRNA (green). rRNA helices h44 and H69 are shown in cartoon in gold and dark blue, respectively. **(C)** Close-up view of Neo1 occupying the conserved pocket formed by h44. **(D)** Close-up view of Neo2 binding pocket formed by residues from h1 and h44. **(E)** Close-up view of Neo3 binding pocket in h28. **(F)** Close-up view of Neo4 binding pocket formed by H69 and stabilized by nucleotides from both H69 and the mtSSU. Key interacting nucleotides are represented as sticks (oxygen in red, nitrogen in blue). **(G)** Conformation-dependent binding patterns of neomycin in different ribosomal rotational states *in vitro*. The dark-yellow dashed semicircle indicates the position of mtSSU in the non-rotated conformation. Solid purple dots represent high binding occupancy, diagonally hatched dots indicate low binding occupancy, and purple open circles (with white lines) denote no observable binding.

Among these, Neo1 and Neo4 correspond to the two canonical aminoglycoside-binding sites previously characterized in bacterial ribosomes. Neo1 displayed well-defined density for all four rings and occupied h44 adjacent to the decoding center A site (**Fig. 1C, Supplementary Figs. S2 and S3A, D**). This position sterically interferes with proper codon–anticodon recognition, consistent with observations in bacterial ribosomes(Wallis et al. 1995, Wang et al. 2012) (**Supplementary Fig. S3A**). Neo1 is coordinated by multiple nucleotides within h44 of the mtSSU (**Fig. 1C, Supplementary Fig. S2D**). Notably, ring I forms a stacking interaction with the nucleobase C1556 (**Fig. 1C, Supplementary Fig. S2D**). In addition, the conserved decoding-center nucleotides A1557 and A1558 (equivalent to A1492/A1493 in bacteria), adopt flipped-out conformations (**Fig. 1C**), characteristic of decoding-center activation and near-cognate tRNA accommodation(Carter et al. 2000, Ogle et al. 2001, Holm et al. 2023). Neo4 binds to the terminal hairpin of H69, representing the second canonical aminoglycoside-binding site (**Fig. 1B, F, Supplementary Fig. S3B**). Although Neo4 exhibits lower occupancy than Neo1, the density is sufficient to assign rings I–III (**Supplementary Fig. S3D**). Neo4 is stabilized by nucleotides from both the mtSSU (G1079 U1081 and G1582) and mtLSU (C2583 and U2586) (**Fig. 1F, Supplementary Fig. S2E**).

Beyond these conserved sites, seven additional neomycin-binding sites are distributed throughout the mitoribosome. Neo2 occupies a pocket formed by h1 and h44 and exhibits well-defined density for all four rings (**Fig. 1D, Supplementary Fig. S3C, D**). Neo3 binds to h28 near the P-tRNA and is stabilized by interactions with surrounding mtSSU nucleotides (**Fig. 1E, Supplementary Fig. S3C, D**). The remaining molecules (Neo5-9) are located at peripheral regions of the ribosome. Together, these observations indicate that neomycin possesses substantial binding landscape across the mitoribosomal rRNA scaffold under *in vitro* conditions.

Analysis of the classified maps reveals conformation-dependent differences in neomycin binding, particularly at the canonical sites h44 (Neo1) and H69 (Neo4) (**Supplementary Table S2**). For Neo1 at h44, density is predominantly observed in non-rotated and partially rotated states, with a weaker but detectable signal in the rotated state (**Supplementary Table S2**). This conformation-dependent pattern is consistent with a preference of Neo1 for non-rotated ribosomal conformations, in line with its inhibitory mechanism in bacterial ribosomes(Wang et al. 2012, Wasserman et al. 2015). Similarly, H69-associated Neo4 density is most apparent in the non-rotated state, where approximately three-ring density can be resolved, and no reliable density is observed in partially rotated or rotated conformations (**Fig. 1F, Supplementary Table S2**). Together, these observations indicate that h44 and H69 preferentially bind neomycin in the non-rotated state, with reduced binding in rotated state (**Fig. 1G**).

### Neomycin exhibits extensive binding despite reduced canonical recognition on the human cytosolic ribosome

To assess the potential impact of neomycin on human cytosolic translation, we first examined its interaction with the cytosolic ribosome. Although sequence divergence within the decoding-center h44 has been proposed to reduce aminoglycoside sensitivity relative to bacterial ribosomes(Fourmy et al. 1996, Borovinskaya et al. 2007, Wang et al. 2012, Wasserman et al. 2015, Paternoga et al. 2023) (**Supplementary Fig. S2A-C**), *in vitro* translation (IVT) assays show that neomycin inhibits translation, with an IC_50_ of approximately 50 μM (**Fig. 2A**). To define the structural basis of this inhibition, we performed cryo-EM analysis of 80S ribosome complexes incubated with 50 μM neomycin (**Supplementary Fig. S4, Supplementary Table S3**). The resulting maps reveal an extensive neomycin-binding landscape consisting of 17 distinct sites, including three on the SSU (Neo1, Neo10 and Neo11) and fourteen on the LSU (Neo12-Neo25) (**Fig. 2B, Supplementary Table S4, Supplementary Fig. S5**).

**Figure 2.**
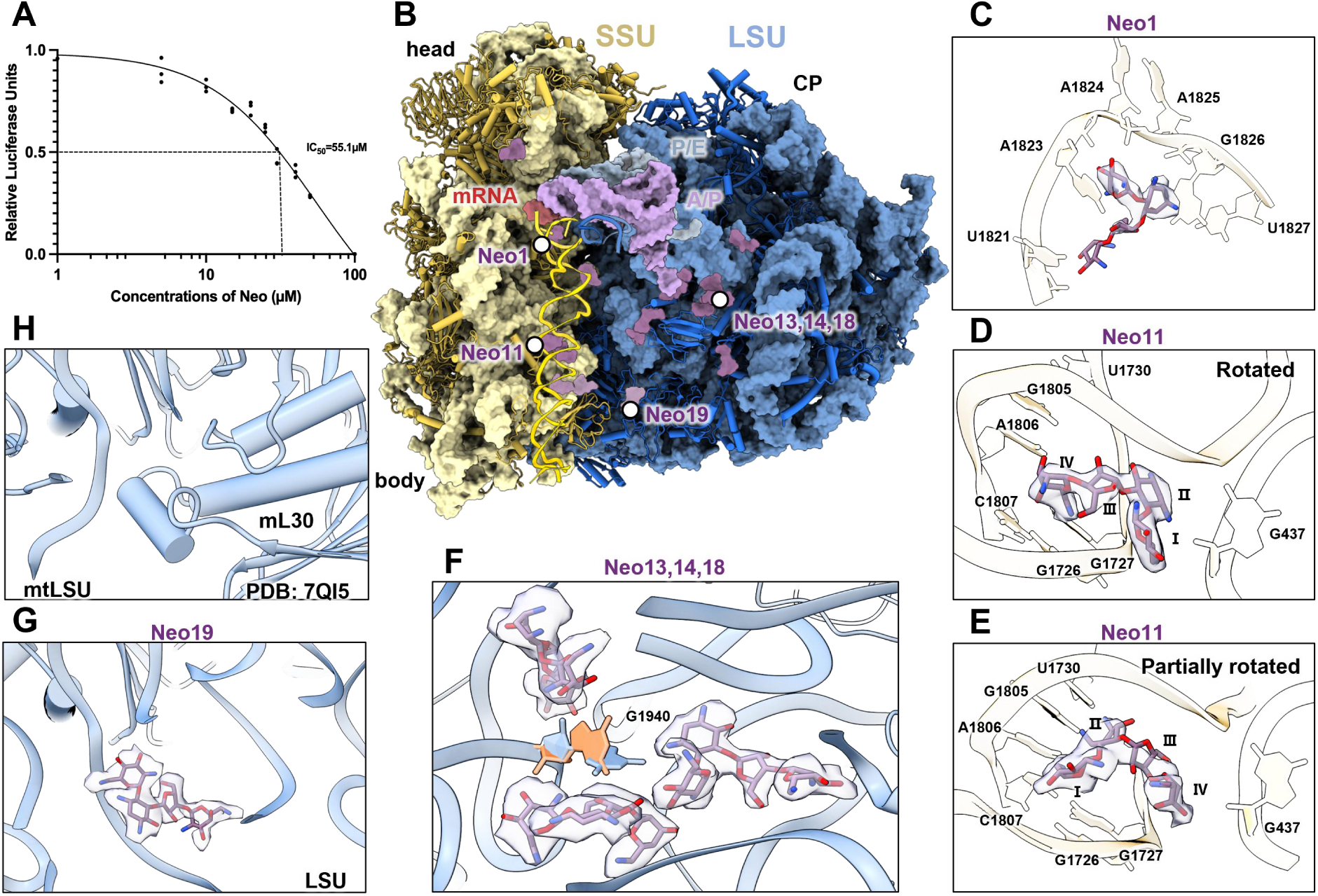
Structural analysis of neomycin binding to cytosolic ribosomes *in vitro*. **(A)** Luciferase reporter assays of cytosolic ribosomes in response to neomycin at different concentrations of 1, 5, 10, 15, 20, 25, 30, 40, 50 and 100 μM (data are presented as mean ± s.d., n = 3 independent experiments). **(B)** Overall structure of neomycin-bound cytosolic ribosomes in rotated state comprising the SSU (yellow), LSU (blue), mRNA (red), A/P-site tRNA (magenta) and P/E-site tRNA (green). rRNA helices h44 and H69 are shown as cartoon in gold and dark blue, respectively. **(C)** Close-up view of Neo1 bound to h44. **(D)** Close-up view of Neo11 bound to a novel pocket within h44 in the rotated state. **(E)** A distinct conformation of Neo11 observed in the partially rotated state. **(F)** A novel binding pocket accommodating three neomycin molecules (Neo13, Neo14, and Neo18), with G1940 adopting a unique conformation upon neomycin binding (orange: pre-binding, blue: post-binding). **(G)** Close-up view of Neo19 bound to a pocket formed by H53 and H54 from the LSU. **(H)** Comparison showing that in the mitoribosome, the Neo19 site is occupied by mitochondrial protein mL30 (PDB: 7QI5(Singh et al. 2024)).

At the decoding center, discernible Neo1 density was detected within h44, indicating that the conserved aminoglycoside-recognition site remains accessible in the human 80S ribosome despite sequence divergence (**Fig. 2C**). However, the corresponding density was markedly weaker than that observed in the 55S, suggesting reduced occupancy or binding stability at this canonical site (**Fig. 2C**). Modeling of the bound neomycin revealed that its presence is associated with the characteristic flipped-out conformations of A1824 and A1825 (A1492/A1493 in bacteria) (**Fig. 2C**), suggesting conservation of decoding-center rearrangements upon aminoglycoside binding.

Structural comparison further reveals subtle alterations in the binding pose relative to bacterial ribosomes, likely arising from local sequence divergence (**Supplementary Fig. S5A-C**). Notably, no neomycin density was observed at the canonical H69 site (**Fig. 2B**). Structural comparison revealed substantial architectural differences in human H69 relative to bacterial ribosomes, resulting in a markedly altered local binding environment (**Supplementary Fig. S5D**). Together, these observations indicate that canonical aminoglycoside recognition is only partially preserved in the human cytosolic ribosome.

Despite this weakened canonical recognition, neomycin engages numerous additional sites throughout the 80S ribosome. Among these, Neo11, located within the central region of h44, exhibits pronounced conformational dependence (**Figs. 2D-E, Supplementary Fig. S5H**). In the rotated state, Neo11 interacts primarily with G437, G1726 and A1806 (**Fig. 2D**). Upon a ∼5° SSU rotation into the partially rotated state, the binding pocket shifts by approximately 3 Å, accompanied by an ∼180° reorientation of the drug molecule (**Fig. 2E**). In this alternate pose, Neo11 establishes a distinct interaction network involving C1807 and U1730 (**Fig. 2E**). These observations indicate that neomycin recognition at non-canonical sites can be sensitive to global ribosome conformation.

Additional structural effects are observed within the LSU. At the H89-H72 junction, the concerted interaction of Neo13, Neo14 and Neo18 is associated with a displacement of nucleotide G1940 (corresponding to bacterial G1026) (**Fig. 2F**). Notably, conformational rearrangements involving the corresponding bacterial G1026 have previously been reported in engineered ribosomes(Huang et al. 2020), raising the possibility that neomycin binding stabilizes alternative conformations within this conserved rRNA region. Furthermore, several neomycin-binding pockets are unique to the human 80S ribosome and occupied regions inaccessible in bacterial or mitochondrial ribosomes due to protein occlusion (**Figs. 2G-H, Supplementary Figs. S5E-G**). These sites further expand the landscape of neomycin-binding interfaces available on the human 80S ribosome.

To compare interaction patterns across binding sites, we quantified atom-level interaction frequencies across previously reported bacterial ribosome structures(Wang et al. 2012, Wasserman et al. 2015) and the human ribosome complexes determined here. Despite substantial variation in binding locations, neomycin interactions were consistently dominated by rings I and II, particularly the amino groups within the conserved 2-DOS core (**Supplementary Fig. S5I**). These recurring interaction modules primarily engage phosphate oxygens and ribose hydroxyl groups within the rRNA backbone, indicating that the chemical basis of neomycin recognition remains highly conserved despite divergence in individual binding pockets.

### Ribosome-associated neomycin is differentially recovered after cellular exposure

Our structural analysis revealed extensive neomycin-binding landscapes on both the mitoribosome and cytosolic ribosome *in vitro*. To assess cellular exposure to neomycin, we first quantified intracellular neomycin accumulation. Using a validated HPLC-UV method following DNS-Cl derivatization, we detected substantial neomycin accumulation within isolated mitochondria following cellular treatment with either 1 mM or 3 mM concentration (**Fig. 3A**). Although direct incubation of isolated mitochondria with 50 µM neomycin, the concentration used for *in vitro* structural assays, yielded the highest accumulation, drug levels achieved through cellular uptake were substantial and within a comparable order of magnitude (**Fig. 3A**). Thus, cellular treatment resulted in substantial mitochondrial exposure to neomycin under the conditions examined.

**Figure 3.**
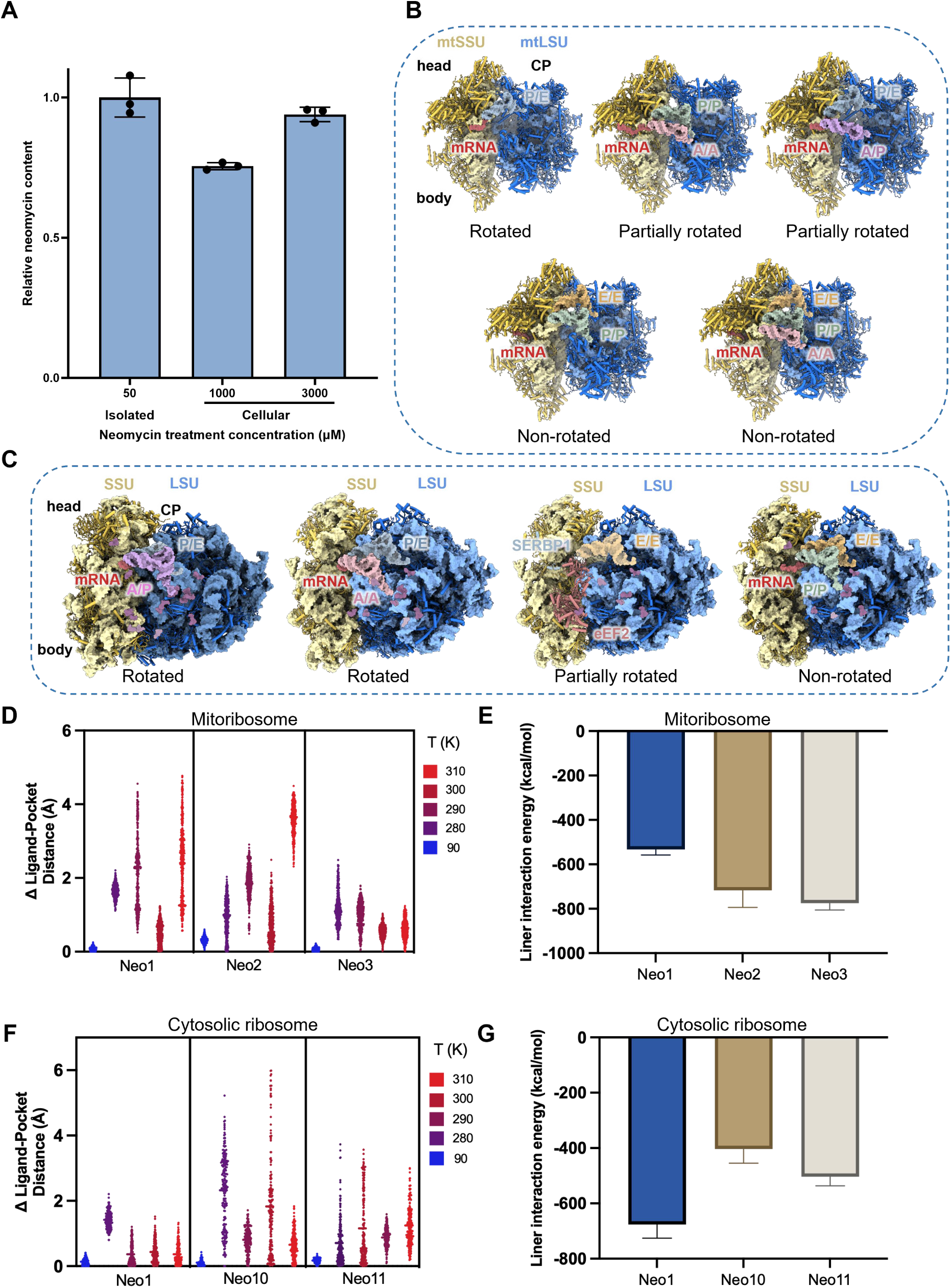
Cellular effects and structural dynamics of neomycin on human ribosomes. **(A)** Quantification of mitochondrial neomycin accumulation. Neomycin levels were quantified by HPLC-UV following pre-column derivatization with dansyl chloride (DNS-Cl). The plot compares mitochondria directly treated *in vitro* (50 µM) with those isolated from cells following 1 h exposure to 1 mM or 3 mM neomycin. Data are presented as relative content normalized to the 50 µM *in vitro* group to facilitate comparison across experimental modes. Data represent mean ± SD (n = 3). **(B)** Overall cryo-EM structure of native mitoribosome with different tRNAs and cofactors. Coloring is consistent with previous figures. **(C)** Overall cryo-EM structure of *in-cell* cytosolic ribosome with different tRNAs and cofactors. Coloring is consistent with previous figures. **(D)** MD simulations depicting the temperature-dependent binding stability of neomycin molecules (Neo1, Neo2, and Neo3) within the mtSSU, illustrating the relative movement of the neomycin molecules within their binding pocket. The changes in distance between the ligand mass center and the binding pocket mass center (defined as the centroid of residues within 3 Å of the ligand in initial cryo-EM structure) during molecular dynamics simulations at temperatures ranging from 90 K to 310 K. **(E)** Calculated linear interaction energy for neomycin within the mtSSU. **(F)** MD simulations depicting the temperature-dependent binding stability of neomycin molecules (Neo1, Neo10, and Neo11) within the cytosolic SSU, illustrating the relative movement of the neomycin molecules within their binding pocket. The changes in distance between the ligand’s center of mass and the binding pocket’s center of mass (defined as the centroid of residues within 3 Å of the ligand in initial cryo-EM structure) during molecular dynamics simulations at temperatures ranging from 90 K to 310 K. **(G)** Calculated linear interaction energy for neomycin within the cytosolic SSU.

To examine whether neomycin-associated density could be detected on mitoribosomes recovered after cellular exposure, we performed cryo-EM analysis of mitoribosomes isolated from HEK293F cells treated with 1 mM neomycin (**Supplementary Fig. S6, Supplementary Table S5**). This concentration is comparable to the millimolar drug accumulation reported in the kidney—a major site of aminoglycoside toxicity(Zaher and Green 2009, Becker and Cooper 2013). Although we minimized the isolation procedure to reduce potential loss of ribosome-associated neomycin, transient interactions may not be retained during isolation. In the mitoribosomes isolated from neomycin-treated cells, no discernible neomycin density was detected, including the canonical aminoglycoside-recognition sites h44 (Neo1) and H69 (Neo4) (**Fig. 3B, Supplementary Fig. S6, Supplementary Table S2**).

We then analyzed cytosolic ribosomes isolated from the same batch of neomycin-treated cells. Structural classification revealed four major translational states, with the rotated state being the most prevalent (**Fig. 3C, Supplementary Fig. S7, Supplementary Table S5**). Neomycin-associated densities remained detectable at most previously assigned sites (14 of 17 sites, **Fig. 3C, Supplementary Table S4**). Notably, the canonical decoding site (Neo1/h44), which displayed relatively weak occupancy even *in vitro*, showed no discernible neomycin density (**Fig. 3C, Supplementary Table S4**). The absence of resolvable neomycin density at this site was not accompanied by detectable structural rearrangements within the h44 pocket (**Supplementary Fig. S8A, B**).

Overall, cellular treatment remodeled the occupancy landscape, with neomycin-associated densities more frequently detected in rotated ribosomal conformations (**Supplementary Table S4**). For example, Neo11, a conformation-dependent binding site within h44, lost density in the non-rotated ribosomes but remained detectable in the rotated conformation (**Supplementary Table S4**). Thus, neomycin-associated densities varied across the different conformational states of the 80S ribosome.

To evaluate whether the altered occupancy patterns could arise from intrinsic instability of the structurally defined binding modes, we performed all-atom explicit-solvent molecular dynamics (MD) simulations of neomycin-bound mitochondrial and cytosolic ribosome complexes. Across a range of temperatures up to physiological conditions (310 K), neomycin remained associated with its corresponding binding pockets despite thermal fluctuations in ligand position and conformation (**Fig. 3D-G, Supplementary Fig. S8C-H**). Consistently, RMSD analysis and linear interaction energy calculations further supported persistent ligand–ribosome interactions over the simulations. These results suggest that the neomycin-binding modes resolved *in vitro* remained stable under physiological temperature conditions.

### Cellular neomycin exposure perturbs translational fidelity without global translation inhibition

We next examined the functional consequences of cellular neomycin exposure for protein synthesis. Neomycin exerted only modest effects on cell viability, with a CCK-8-derived IC_50_ of approximately 1.2 mM after 72 h exposure (**Fig. 4A**). Similarly, L-azidohomoalanine (L-AHA) labeling revealed no significant reduction in either mitochondrial or cytosolic protein synthesis following neomycin treatment (**Fig. 4B-C**), and only a modest reduction in nanoluciferase reporter expression (**Fig. 4D, Supplementary Fig. S9A, C-G**). Together, these results indicate that neomycin does not cause widespread translational shutdown under the cellular conditions examined.

**Figure 4.**
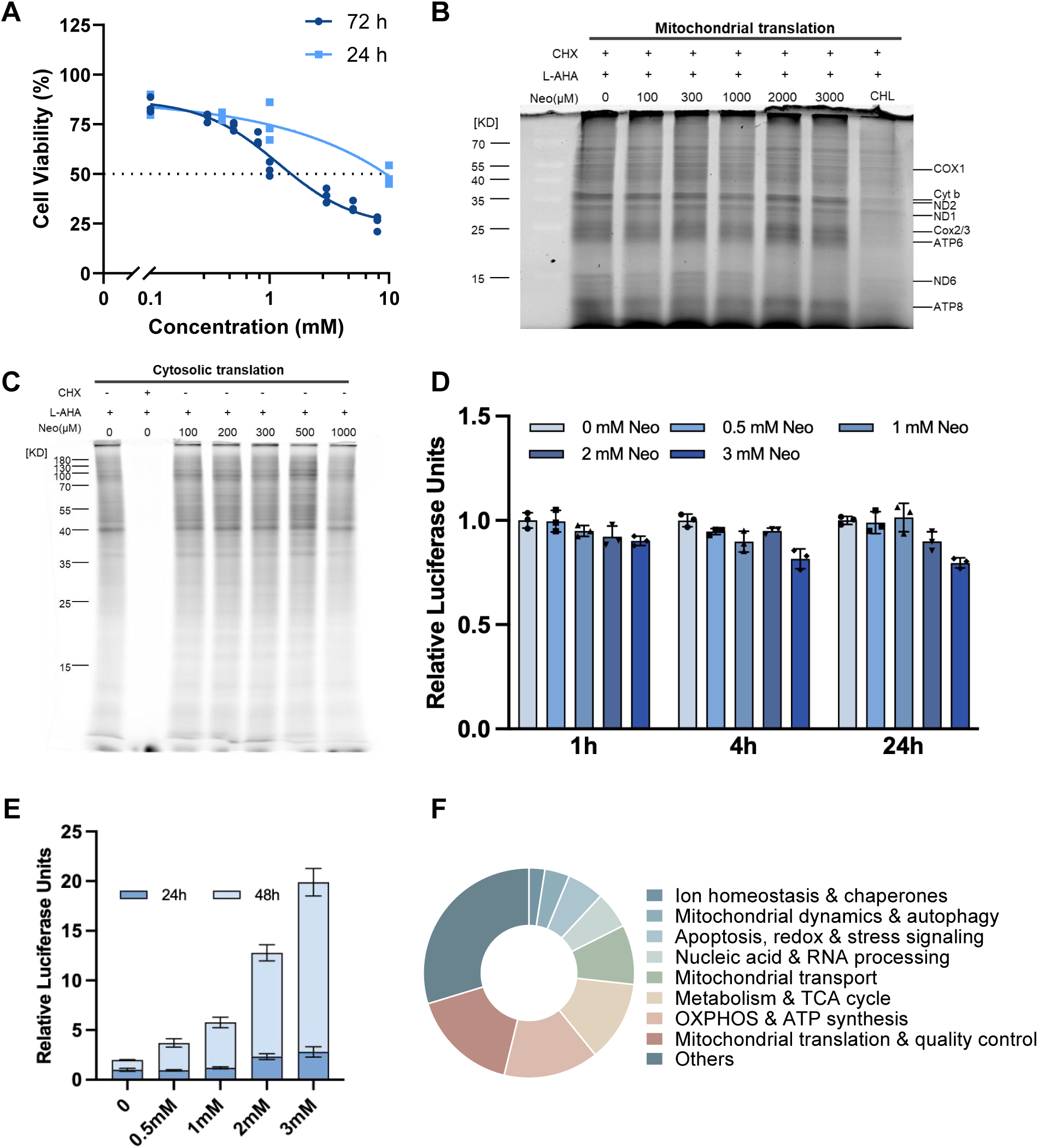
Cellular effects of neomycin with human ribosomes. **(A)** Cell viability of HEK293T cells measured by the CCK-8 assay after treatment with increasing concentrations of neomycin for 24 h and 72 h, respectively (data are presented as mean ± s.d., n = 3 independent experiments). **(B)** *De novo* mitochondrial nascent protein synthesis measured via L-AHA labeling. CHL, chloramphenicol. **(C)** *De novo* cytosolic nascent protein synthesis measured via L-AHA labeling. **(D)** Cellular translation assays using the NLuc cell line at various time points following treatment with different concentrations of neomycin (data are presented as mean ± s.d., n = 3 independent experiments). **(E)** Quantification of cystic fibrosis transmembrane conductance regulator (CFTR) nonsense allele readthrough in cultured HEK293T cells treated with increasing concentrations of neomycin for 24 h and 48 h (data are presented as mean ± s.d., n = 3 independent experiments). **(F)** Functional classification of mitochondria-related genes exhibiting neomycin-induced readthrough. Donut chart showing the distribution of 370 mitochondria-related genes exhibiting significant stop-codon readthrough induction. Genes are clustered into essential pathways including OXPHOS assembly, mitochondrial translation, and protein quality control (PQC)/stress response.

Although neomycin had minimal effects on global protein synthesis, we observed a concentration-dependent increase in stop-codon readthrough in a reporter cell line upon neomycin treatment, indicating a selective effect on translational fidelity (**Fig. 4E, Supplementary Fig. S9B**). To determine whether this effect extended beyond reporter-based measurements, we analyzed a published genome-wide Ribo-seq dataset obtained following neomycin treatment(Wangen and Green 2020). Consistent with our cellular assays, global translation efficiency remained largely unchanged(Wangen and Green 2020). In contrast, the Ribosome Readthrough Score (RRTS) was significantly elevated across the transcriptome, indicating widespread translational readthrough (**Supplementary Fig. S10A**). Among 2,102 nuclear-encoded transcripts exhibiting significantly increased readthrough, 370 were associated with mitochondrial pathways, including respiratory chain assembly, organellar translation, and stress response (**Fig. 4F, Supplementary Fig. S10B**).

Together, our structural and functional analyses reveal extensive neomycin binding to human ribosomes *in vitro*, while cellular exposure leads to altered translational fidelity without widespread inhibition of protein synthesis (**Fig. 5**).

**Figure 5.**
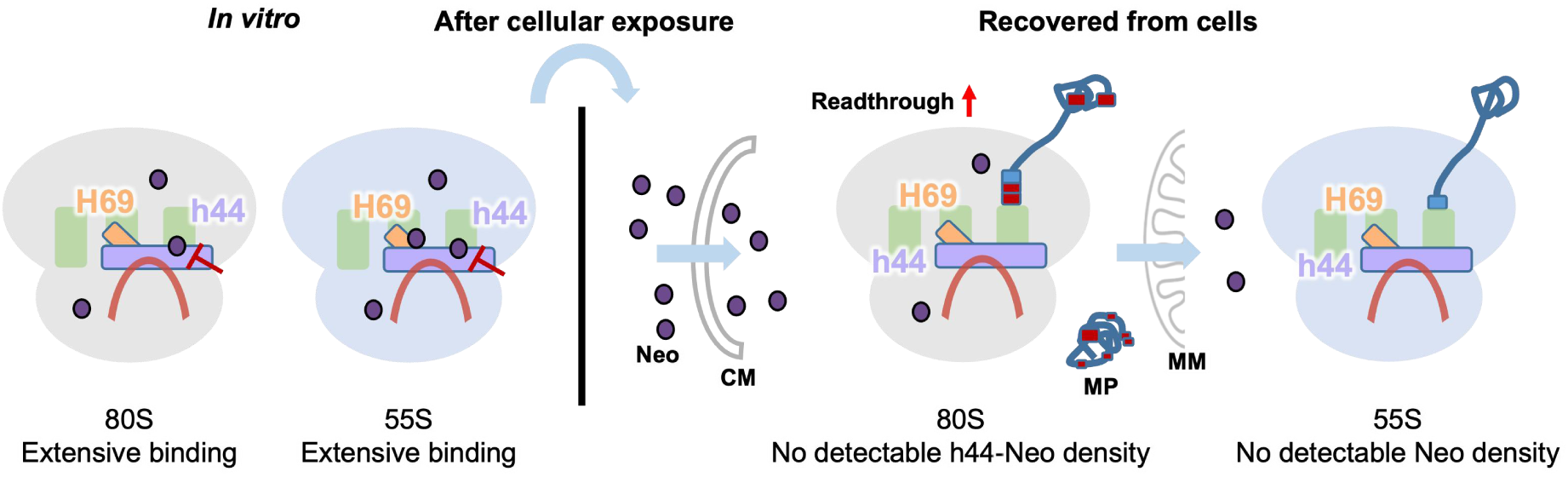
Divergence between structural binding landscape and cellularly recovered ribosome-associated neomycin states. Left (*in vitro*): Isolated 80S and 55S ribosomes show extensive multi-site neomycin occupancy (purple spheres), revealing the structural binding landscape of neomycin under defined biochemical conditions. Right (ribosomes recovered from neomycin-treated cells): 80S ribosomes recovered from treated cells show no detectable h44-associated neomycin density, whereas neomycin treatment induces increased stop-codon readthrough at the cellular level. Similarly, 55S ribosomes recovered from treated cells show no detectable neomycin density, despite confirmed mitochondrial drug accumulation. CM, cell membrane; MM, mitochondrial membrane; MP, misfolded protein.

## Discussion

Our study provides a structural and cellular analysis of neomycin interaction with human ribosomes, defining a broad binding landscape across both mitochondrial and cytosolic ribosomes *in vitro*. Neomycin binds at multiple sites on both ribosomes, including conserved regions of the decoding center despite differences in rRNA sequence. Following cellular exposure of neomycin, ribosomes isolated from treated cells showed different patterns of associated neomycin, with no detectable density on the mitoribosomes and reduced occupancy at a subset of cytosolic ribosome sites. These observations indicate that the broad binding landscape defined *in vitro* is not equivalently represented among drug-associated densities recovered after cellular exposure.

Several factors may contribute to this difference. Unlike the simplified biochemical environments used for structural characterization, the intracellular environment is characterized by high ionic strength, macromolecular crowding, and competitive binding from endogenous polyamines and tRNAs(Cortese et al. 1991, Prezant et al. 1993, Marín et al. 1995, Ellis 2001, Spitzer and Poolman 2009). These conditions may alter the stability and dynamics of drug–ribosome interactions, a phenomenon consistent with emerging evidence that the intracellular environment critically influences the efficacy of aminoglycosides(Dey et al. 2025). In addition, weak or transient interactions may dissociate during ribosome isolation. Although our MD simulations support the local stability of structure-resolved neomycin-binding poses under physiological temperature conditions, they do not capture the long-timescale dissociation kinetics that may occur during ribosome purification. Thus, the ribosome-associated neomycin densities likely represent the subset of interactions sufficiently stable to withstand the isolation procedures.

Furthermore, cellular exposure to neomycin revealed a distinct functional response. Despite minimal effects on global protein synthesis, neomycin consistently promotes translational readthrough in both reporter assays and transcriptome-wide analyses. This is consistent with the possibility that dynamic interactions at functionally relevant ribosomal sites, including the decoding center, may contribute to altered translational fidelity(Kolosova et al. 2025). Notably, proteins associated with mitochondrial functions were among those affected by translational readthrough, suggesting that altered translational fidelity may have broader cellular consequences beyond the global translation output.

Aminoglycoside-associated toxicity has been linked to mitochondrial translation defects, particularly in individuals carrying mitochondrial rRNA variants such as A1555G. Previous studies have suggested that A1555G shifts the mitochondrial decoding-center architecture toward a more bacterial-like configuration, providing a possible structural explanation for increased aminoglycoside susceptibility(Prezant et al. 1993, Guan 2011, Greber et al. 2015). While we did not resolve neomycin density on 55S ribosomes from treated cells, this finding does not exclude mitochondrial contributions to neomycin-associated cellular effects, particularly in genetically susceptible contexts. Together with the altered translational fidelity observed in cells, these findings support a multifaceted view of aminoglycoside action, in which mitochondrial ribosome susceptibility and translational fidelity may contribute to cellular responses in a context-dependent manner.

Overall, our work provides a comprehensive analysis of neomycin interactions with human mitochondrial and cytosolic ribosomes. By defining a broad structural binding landscape and integrating these findings with cellular functional consequences, our study provides a framework for understanding aminoglycoside–ribosome interactions in human cells. More broadly, our study highlights the importance of considering native cellular environments when evaluating the efficacy and toxicity of RNA-targeting compounds.

## MATERIAL AND METHODS

### Cell lines and cell culture

HEK293T cells were cultured at 37 °C and 5% CO_2_ in Dulbecco’s Modified Eagle’s Medium (DMEM, Gibco, C11995500BT) supplemented with 10% fetal bovine serum (FBS, Excell Bio, FSP500). HEK293F cells were cultured in SMM 293-TII expression medium (Sino Bio, M293TII) in shake flasks at 37 °C, 5% CO_2_ and 110 rpm.

### Plasmid construction and generation of stable cell lines

Clonal stable cell lines expressing NanoLuciferase (NLuc) or cystic fibrosis transmembrane conductance regulator (CFTR) were generated as previously described(Li et al. 2020). The NLuc expression construct was generated by PCR amplification of NLuc-Flag fragments, subsequently assembled into a linearized vector (originally derived from Jamie H. D. Cate’s laboratory) via Gibson Assembly. For the *CFTR* plasmid, *CFTR-G542X (UGA)* along with the G542X context (seven codons upstream and downstream) was fused to NLuc. This recombinant fragment was cloned into the same laboratory-derived backbone vector using Gibson Assembly. The CFTR-G542X mutant was created through site-directed mutagenesis of wild-type *CFTR* cDNA templates. The plasmid construct was subsequently transfected into HEK293T cells and stable transfectants were selected using 1 μg/ml puromycin for two weeks.

### Cell-based translation assays

The effect of neomycin on cytosolic translation was evaluated using the NLuc cell lines, which were exposed to different concentrations of neomycin (0.5, 1, 2 and 3 mM) or DMEM as a control. Luciferase activity was quantified after 1, 4, and 24 h using microplate luminometer (BERTHOLD, Centro XS3 LB 960). To assess the impact of the redox environment on neomycin toxicity, 0.3 mM H_2_O_2_ was co-administered with varying concentrations of neomycin (0.1, 1, and 3 mM) for 1 h. Additionally, to determine the readthrough effect of neomycin, the CFTR cell lines were treated with different concentrations of neomycin (0.5, 1, 2, and 3 mM) for 24 h and 48 h, respectively.

### *In vitro* transcription and translation assays

The DNA template for *in vitro* transcription was PCR-amplified from plasmid pGM_EMCV-NLuc using primers encoding T7 RNA polymerase promoter and poly-A tail sequences. The *in vitro* transcription reaction was performed as previously described(Li et al. 2020). mRNA transcripts were assessed for quality and size by gel electrophoresis, quantified spectrophotometrically (Implen spectrophotometer, Nanophotometer NP80), diluted to 1 μg/µL, aliquoted, and preserved at -80 °C prior to *in vitro* translation reactions.

HEK293T cells were harvested at ∼80% confluency and resuspended in ice-cold lysis buffer (10 mM HEPES-KOH pH 7.5, 10 mM KOAc, 0.5 mM Mg(OAc)_2_, 5 mM DTT) at a 1:1 (v/v) ratio. The HEK293T cell lysates and IVT reaction systems were prepared as described previously(Lintner et al. 2017, Li et al. 2019). Neomycin (MCE, HY-17624A) was introduced at different concentrations as experimental variables in the reaction system at final concentrations of 1 µM, 5 µM, 10 µM, 15 µM, 25 µM, 30 µM, 40 µM, 50 µM and 100 µM. Reactions were equilibrated at 30 °C for 25 min prior to luciferase activity measurement using a microplate luminometer (BERTHOLD, Centro XS3 LB 960). Lysates and mRNA aliquots from identical preparation lots were used for comparative assays within individual experimental series to control batch variability.

### Cell counting kit-8 (CCK-8) cell growth assay

The cytotoxicity of neomycin was measured in HEK293T cell lines using the CCK-8 assay. Approximately 8000 cells per well were seeded in a 96-well plate in DMEM medium supplemented with 10% FBS. After 24 h of adhesion, the cells were treated with different concentrations of neomycin dilutions versus DMEM vehicle control. After 72 h of incubation, culture medium was replaced with fresh DMEM medium supplemented with CCK-8 (10% v/v reagent: medium ratio) for an additional 2 h of light-protected incubation. Cellular viability was determined by measuring at 450 nm using a microplate reader. The half-maximal inhibitory concentration (IC_50_) for neomycin was calculated by fitting the dose-response curve using GraphPad Prism software. All experiments were performed in biological triplicates (n = 3).

### L-AHA labeling of nascent polypeptides

L-AHA labeling of mitochondrial-encoded nascent polypeptides was performed as described previously(Li et al. 2024). Briefly, the cells were treated with different concentrations of neomycin and 50 μg/mL cycloheximide (CHX, MCE, HY-12320), which selectively inhibits eukaryotic cytosolic ribosome activity by blocking translational elongation. Fluorescence signals were analyzed using an Amersham Typhoon 5 (Cytiva).

For the cytosolic nascent polypeptide labeling, the procedure mirrored that used for mitochondrial-encoded nascent polypeptides, with cells treated with 50 μg/mL CHX as a positive control. Following the labeling, cells were harvested and resuspended in buffer containing 60 mM Tris-HCl pH 8.0, 50 mM NaCl, 2 mM MgCl_2_, 40 mM KCl, 5% glycerol, 0.1% NP-40 and 1× protease inhibitor and were subsequently homogenized on ice using a glass dounce homogenizer. The resulting cell lysate was cleared by centrifugation at 15,000 *×g* for 15 min at 4 °C, and the supernatant was transferred to new 1.5 mL tubes. The click reaction and the following protein signal detection were performed as described above. All experiments were performed in triplicate (n = 3 independent experiments).

### Isolation of mitochondria and purification of mitoribosomes

Mitochondria isolation was performed as described previously(Aibara et al. 2020). A final volume of 1L HEK293F cells at a density of 3× 10^6^ cells/ml was harvested, washed with Phosphate Buffered Saline (PBS) buffer, and resuspended in 60mL mitochondrial isolation buffer (50 mM HEPES-KOH pH 7.5, 10 mM KCl, 1.5 mM MgCl_2_, 1 mM EDTA, 1 mM EGTA, 1 mM DTT, protease inhibitors). Cells were lysed by grinding in a glass dounce homogenizer and lysate was clarified by centrifugation at 1000 *×g* for 15 min, then 10,000 *×g* for 15 min at 4°C to get the crude mitochondria in the pellet. The resulting pellet was resuspended in 10 mL resuspension buffer (50 mM HEPES-KOH pH 7.5, 10 mM KCl, 1.5 mM MgCl_2_, 1 mM EDTA, 1 mM EGTA, 1 mM DTT, 70 mM sucrose, 210 mM mannitol, and 1× protease inhibitor). Genomic DNA was removed by addition of 100 U RNase-free DNase I and centrifugation at 10,000 *×g* for 15 min at 4 °C. The crude mitochondria were gently resuspended in 1 mL buffer containing 20 mM HEPES-KOH pH 7.5, 1 mM EDTA, 250 mM sucrose and loaded onto sucrose gradients comprising 1.5 mL 60% (w/v), 4 mL 32% (w/v), 1.5 mL 23% (w/v) and 1.5 mL 15% (w/v) sucrose solutions in 20 mM HEPES-KOH pH 7.5, 1 mM EDTA. The gradients were centrifuged at 28,000 rpm for 1h at 4 °C using an SW41Ti rotor (Beckman Coulter). The brown band containing mitochondria migrating to the interface of 32% and 60% sucrose was collected.

The fresh isolated mitochondria were resuspended in lysis buffer (25 mM HEPES-KOH pH 7.5, 100 mM KCl, 10 mM Mg(OAc)_2_, 1.7% Triton X-100, 2 mM DTT, 1× protease inhibitor) at a mitochondria-to-buffer ratio of 1:2 (v/v). The mixture was homogenized with a glass dounce homogenizer. The lysate was cleared by centrifugation at 30,000 *×g* for 20 min at 4 °C. The supernatant was then loaded onto the top of sucrose cushion (1 M sucrose in 20 mM HEPES-KOH pH 7.5, 100 mM KCl, 20 mM Mg(OAc)_2_, 1% TX-100, 2 mM DTT) and centrifuged at 339,200 *×g* for 45 min at 4°C using an MLA130 rotor (Beckman Coulter). The pellet was rinsed and resuspended in Grid buffer (20 mM HEPES-KOH pH 7.5, 100 mM KCl, 10 mM Mg(OAc)_2_, 0.02% DDM, 1 mM DTT). Mitoribosomes (280 nM) were incubated with 50 µM neomycin (∼178-fold excess) for 30 min at 4 °C prior to the cryo-EM sample preparation (molar ratio of ribosome to neomycin around 1:170).

### Analysis of mitochondrial neomycin via HPLC with pre-column derivatization

For the quantification of intramitochondrial neomycin, HEK293T cells were cultured in DMEM supplemented with 10% FBS and exposed to 1 mM or 3 mM neomycin for 1 hour. An in-cell control group was prepared by incubating mitochondria isolated from untreated cells with 50 µM neomycin for 30 min on ice. Mitochondrial fractions from all groups were isolated using differential centrifugation as described previously, and total protein concentrations were determined using the BCA assay to ensure consistent normalization for downstream analysis.

The intramitochondrial neomycin content from the three distinct sample types was quantified using a validated HPLC-UV method following dansyl chloride (DNS-Cl) derivatization. Isolated mitochondrial pellets were resuspended in ice-cold lysis buffer consisting of PBS at pH 7.4 containing 0.5% Triton X-100 and subjected to three freeze-thaw cycles using liquid nitrogen and a 37°C water bath combined with sonication to ensure complete membrane disruption. Proteins were subsequently precipitated by adding an equal volume of ice-cold acetonitrile and the supernatant was clarified by centrifugation at 15,000 rpm for 30 min at 4°C. To recover low-molecular-weight analytes including neomycin, the supernatant was transferred to a 3 kDa molecular weight cut-off (MWCO) ultrafiltration device and centrifuged to obtain the final extract.

The extracted samples or neomycin reference standards were derivatized by the sequential addition of an equal volume of 0.2 M sodium bicarbonate buffer at pH 9.8 and an equal volume of DNS-Cl reagent at a concentration of 2 mg/mL in acetonitrile. The reaction proceeded at 60°C for 40 min in the dark and was terminated with the addition of 1% acetic acid. The derivatized samples were analyzed using a reversed-phase C18 column on an Agilent 1260 Infinity II HPLC system. Chromatographic separation was achieved with a water-acetonitrile gradient at a flow rate of 0.6 mL/min and UV detection was performed at a wavelength of 250 nm. Neomycin concentrations in the mitochondrial samples were determined from a standard curve generated from neomycin reference standards processed identically to the biological samples and the final values were normalized to the total mitochondrial protein weight.

### Human cytosolic ribosome purification

The human cytosolic ribosome was isolated using a modified protocol adapted from a previously published paper(Holvec et al. 2024). HEK293T cells were harvested at ∼80% confluency, centrifuged at 1000 *×g* for 10 min at 4 °C, and washed twice with ice-cold PBS. The cell pellet was resuspended in an equal volume of lysis buffer (10 mM HEPES-KOH pH 7.5, 10 mM KOAc, 0.5 mM Mg(OAc)_2_, 5 mM DTT). Following 20 min incubation on ice, the resuspended cells were mechanically lysed by grinding in a glass homogenizer. Lysates were centrifuged at 1,000 *×g* for 10 min to remove cell debris, and cleared by centrifugation at 10,000 *×g* for 15 min at 4 °C. The supernatant was then loaded onto a sucrose cushion (10 mM HEPES-KOH pH 7.5, 50 mM KCl, 5mM Mg(OAc)_2_, 0.02% DDM, 1 mM DTT, 1M sucrose) and centrifuged at 603,000 *×g* for 45 min at 4 °C in an MLA130 rotor (Beckman Coulter). The pellets obtained were washed and sequentially resuspended in Grid buffer (20 mM HEPES-KOH pH 7.5, 50 mM KCl, 5 mM Mg(OAc)_2_, 0.02% DDM) and measured using an Implen spectrophotometer. Cytosolic ribosomes (294 nM) were incubated with 50 µM neomycin (∼170-fold excess) for 30 min at 4 °C prior to the cryo-EM sample preparation (molar ratio of ribosome to neomycin around 1:170).

### Isolation of mitoribosome and cytosolic ribosome from neomycin treated cells

HEK293F cells were scaled up sequentially, by inoculating at 1×10^6^ cells/mL and subsequently splitting at a cell density of 3.0×10^6^ cells/mL until achieving 300 mL final culture volume. Neomycin was added to HEK293F cells to a final concentration of 1 mM for 1 hour prior to utilization. Native human mitoribosomes purification was performed as described for the “Isolation of mitochondria and purification of mitoribosomes” part. And no extra neomycin was added during the purification. The same batch of cells were used for cytosolic ribosome isolation to make strict comparison, meaning we have native mitoribosomes and cytosolic ribosomes from the same cells. After cells harvest and lysis, the supernatant was applied for cytosolic ribosome isolation and the pellet was used for crude mitochondria isolation. The protocol for purification of cytosolic ribosomes is the same as described in “Human cytosolic ribosome purification”.

### Cryo-EM grids preparation

The concentrations of the mitoribosomes and cytosolic ribosomes used for cryo-EM grid preparation were 280 nM and 160 nM, respectively. Approximately, 3.2 μL aliquots were applied to a plasma-cleaned holey carbon grid (R1.2/1.3, 300 mesh, Quantfoil) for 1 min, on which a homemade continuous carbon film was precoated. Grids were blotted for 2.5 s under 100% humidity at 4 °C and plunge-frozen in liquid ethane using Vitrobot Mark IV (Thermo Fisher Scientific).

### Cryo-EM data collection and processing

Cryo-EM micrographs of all samples were collected on a Titan Krios transmission electron microscope (Thermo Fisher Scientific) operated at 300 kV. The micrographs were recorded using a Falcon 4i direct electron detector (Thermo Fisher Scientific) at a pixel size of 0.81 Å. A total electron dose of 50 electrons per Å² was fractionated over 40 frames with a total exposure time of 5.9 s. Data were acquired using EPU software(Thompson et al. 2019) with a defocus ranging from 1.0 to 2.0 μm.

All datasets were processed using cryoSPARC v.4.2.1(Punjani et al. 2017). The movie stacks were first corrected for motion using the patch motion correction tool of cryoSPARC. Contrast transfer function (CTF) parameters were estimated using the patch CTF estimation tool. Manual curation was performed to exclude suboptimal micrographs. From the curated micrographs, particles were initially identified through blob-based template-free picking, followed by iterative template-based particle picking using averaging projections selected from 2D classification. The extracted particles underwent multiple rounds of 2D classification to remove non-ribosomal contaminants and ice artifacts. Particles were then used for 3D auto-refinement yielding a map for further classification.

Focused 3D classification was carried out using a mask on the SSU to facilitate conformational sorting, enabling separation of rotated versus non-rotated ribosome states. Another round of focused 3D classification using a spherical mask on the tRNA binding sites was performed, separating the particles into different tRNA binding states. H69 and h44 are key binding sites of aminoglycoside antibiotics. Therefore, a spherical mask of radius 23 Å was generated for H69 and h44, respectively, for 3D classification to improve the local resolution. Masks with smoothed edges were generated with the "relion_mask_create" tool in RELION4.0.0(Kimanius et al. 2021). After one round of CTF refinement, particles of each class were further processed by focused refinements with different masks applied. All reported map resolutions were determined by the FSC = 0.143 criterion.

### Model building and refinement

In general, the structures of cytosolic ribosomes (PDB: 8QOI(Holvec et al. 2024)) and mitoribosome (PDB: 7QI5 and 7QI6(Singh et al. 2024)) were used for rigid body docking into the cryo-EM density map using UCSF ChimeraX v1.8(Goddard et al. 2018). The docked coordinates were realigned to match map orientation and saved as models for manual building and adjustment in Coot v0.9.8(Emsley et al. 2010).

Extra densities that could not be assigned to ribosomal components were systematically inspected along the rRNA chains. Candidate neomycin densities were evaluated based on their contour level, size, shape, local structural environment, and compatibility with the branched four-ring architecture of neomycin. A contour level of 3σ was used as the primary criterion for density assignment, while maps were additionally inspected across a broader contour range (1.8–3.5σ for cytosolic ribosomes and 2.4–3.5σ for mitoribosomes) to assess the consistency of candidate densities. Densities compatible with the size and shape of neomycin were distinguished from small spherical densities consistent with inorganic ions such as Mg²⁺ and elongated densities associated with endogenous polyamines.

The neomycin ligand was fetched from the Coot ligand library using the three-letter code: NMY and manually positioned according to the density maps. The fitted conformations were further adjusted and refined by Coot real-space refinement with appropriate geometric restraints. For the canonical decoding-center binding site, the modeled neomycin conformation was compared with previously reported aminoglycoside–ribosome structures. Conformational changes induced by neomycin binding were also manually adjusted.

Unknown continuous densities were analyzed using CryoAtom(Su et al. 2025) to predict the sequence and three-dimensional structure, which was subsequently added into the model. Final models were further subjected to refinement with Phenix.real_space_refine v1.13_2998(Afonine et al. 2018, Liebschner et al. 2019). The figures were prepared with ChimeraX v1.8(Goddard et al. 2018). Figures were compiled using Adobe Illustrator.

### Computational methods

mtSSU molecular dynamics (MD) simulations were performed using the AMBER package. The system comprised ribosomal proteins, rRNA, and three neomycin molecules (Neo1, Neo2, Neo3) solvated in an orthorhombic TIP3P water box extending 12 Å beyond the solute. The ff19SB force field was applied to proteins, OL3 to RNA, and GAFF2 parameters with AM1-BCC charges to ligands. After energy minimization and stepwise heating, production runs were conducted at 90 K, 280 K, 290 K, 300 K, and 310 K under NPT conditions using a 2-fs timestep with SHAKE constraints. Five independent 100-ns replicates per temperature were acquired. Binding free energies were calculated via the linear interaction energy method using the *cpptraj* module in Ambertools. And the MD of the SSU of the cytosolic ribosome with three neomycin molecules (Neo1, Neo10, and Neo11) were performed as described for the mtSSU.

### Neomycin occupancy estimation

Neomycin occupancy was estimated for both the mitoribosomes and cytosolic ribosomes using OccuPy (v0.1.9)(Forsberg et al. 2023), following the software’s documentation for compositional heterogeneity analysis. Briefly, we processed the input cryo-EM maps through a local filtering approach to generate local resolution maps. These maps were subsequently used to modify the original density maps by applying an amplification power and a final output low-pass filter. The final neomycin occupancy was visualized by coloring the original density maps according to the modified output maps in ChimeraX v1.8(Goddard et al. 2018). This process effectively highlighted the regions of neomycin binding across both ribosome types.

### Ribo-seq data re-analysis and readthrough quantification

To evaluate neomycin-induced translational fidelity defects, published Ribo-seq data from 293T cells (2 mg/mL neomycin, 24 h; GSE138643)(Wangen and Green 2020) were re-analyzed. Sequencing reads were processed and aligned to the human genome as previously described. Metagene plots surrounding canonical stop codons were generated to visualize ribosome density transitions in the 3′UTR. Translational readthrough was quantified by the Ribosome Readthrough Score (RRTS), defined as the ratio of normalized footprint density in the 3′ UTR (between the normal termination codon and the first in-frame stop codon in 3′UTR) versus the CDS. Significant readthrough induction was defined by: (i) *de novo* detection (RRTS_untr = 0 and RRTS_neo > 0); or (ii) at least a two-fold RRTS increase (log_2_(RRTS_neo / RRTS_ctrl) > 1).

Functional annotation of the 2,102 identified candidates (nuclear-encoded transcripts) was performed using DAVID(Huang et al. 2009, Sherman et al. 2022). Within this group, 370 mitochondria-related genes were prioritized and categorized into functional modules—including OXPHOS assembly, organellar translation, and protein quality control (PQC)—to assess the systemic impact on mitochondrial proteostasis. Proportional distributions were visualized via donut charts.

## ACKNOWLEDGEMENTS

We thank Zhengkun Xie from the Center of Advanced Analysis & Gene Sequencing of Zhengzhou University for technical support, Alexey Amunts for discussions and comments on the manuscript, and Jamie Doudna Cate for discussions and suggestions.

## AUTHOR CONTRIBUTIONS

X.Z., L.Z., C.J. and M.W. performed the sample preparation, acquired cryo-EM data, and carried out image processing and structure refinement. L.Z., C.J., M.W. and W.L. did the model building and structural analysis. X.Z., L.Z. and K.W. conducted biochemistry and cell-based experiments. X.H. performed the MD experiment. L.Z., W.L., C.J., and M.W. prepared the figures. W.L. wrote the initial draft of the paper. All authors analyzed the data and edited the paper. W.L., X.H., Z.Z. and L.G. jointly supervised this work and are corresponding authors.

## SUPPLEMENTARY DATA

Supplementary Data are available at NAR online.

## DATA AVAILABILITY

All cryo-EM maps and atomic coordinates generated in this study have been deposited in the Electron Microscopy Data Bank and the Protein Data Bank under the following accession numbers: EMD-65426 and 9W80 (*in vitro* dataset, mitoribosomes in non-rotated state with A/A P/P E/E-site tRNA), EMD-65433 and 9VXM (*in vitro* dataset, mitoribosomes in non-rotated state with A/A P/P-site tRNA), EMD-65428 and 9W85 (*in vitro* dataset, mitoribosome in partially rotated state with A/P P/E-site tRNA), EMD-65427 and 9W86 (*in vitro* dataset, mitoribosome in partially rotated state with A/A P/P E/E-site tRNA), EMD-65431 and 9W84 (*in vitro* dataset, mitoribosome in rotated state with A/P P/E-site tRNA), EMD-65429 and 9W82 (*in vitro* dataset, mitoribosome in rotated); EMD-65419 and 9W7Z (*in vitro* dataset, cytosolic ribosome in non-rotated state with P/P E/E-site tRNA), EMD-65412 and 9W83 (*in vitro* dataset, cytosolic ribosome in partially rotated state with eEF2, SERBP1 and E/E-site tRNA), EMD-65434 and 9VXN (*in vitro* dataset, cytosolic ribosome in rotated state with A/P P/E-site tRNA), EMD-65411 (1 mM native dataset, cytosolic ribosome in non-rotated state with P/P E/E-site tRNA), EMD-65413 (1 mM native dataset, cytosolic ribosome in partially-rotated state with eEF2, SERBP1 and E/E-site tRNA), EMD-65415 (1 mM native dataset, cytosolic ribosome in rotated state with A/P P/E-site tRNA), EMD-65414 (1 mM native dataset, cytosolic ribosome in rotated state with A/A P/E-site tRNA); EMD-65421 (1 mM native dataset, mitoribosome in non-rotated state with P/P E/E-site tRNA), EMD-65422 (1 mM native dataset, mitoribosome in non-rotated state with A/A P/P E/E-site tRNA), EMD-65423 (1 mM native dataset, mitoribosome in partially rotated state with A/A P/P-site tRNA); EMD-65424 (1 mM native dataset, mitoribosome in partially rotated state with A/P P/E-site tRNA), EMD-65425 (1 mM native dataset, mitoribosome in rotated state with P/E-site tRNA), EMD-65420 (1 mM native dataset, LSU of the mitoribosome). Source data are provided with this paper.

The raw ribosome profiling sequencing data analyzed in this study were obtained from the NCBI Gene Expression Omnibus (GEO) under accession number GSE138643 (https://www.ncbi.nlm.nih.gov/geo/query/acc.cgi?acc=GSE138643). These data were deposited in GEO in 2019 and were originally published and described in a study by Wangen JR and Green R (2020) "Stop Codon Context Influences Genome-Wide Stimulation of Termination Codon Readthrough by Aminoglycosides." (DOI: https://doi.org/10.7554/eLife.52611). Source data from ribosome profiling analysis used for Gene Ontology (GO) enrichment analysis are publicly accessible through the *eLife* article page associated with the DOI above (see the file elife-52611-fig2-data1-v2.xlsx in the article’s source data). The GO enrichment analysis data generated by this study are available in the Source Data file of this article.

## REFERENCES

Afonine, P. V., B. K. Poon, R. J. Read, O. V. Sobolev, T. C. Terwilliger, A. Urzhumtsev, and P. D. Adams. 2018. Real-space refinement in PHENIX for cryo-EM and crystallography. Acta crystallographica. Section D, Structural biology 74:531–544.

Aibara, S., V. Singh, A. Modelska, and A. Amunts. 2020. Structural basis of mitochondrial translation. eLife 9.

Amunts, A., A. Brown, J. Toots, S. H. W. Scheres, and V. Ramakrishnan. 2015. Ribosome. The structure of the human mitochondrial ribosome. Science (New York, N.Y.) 348:95–98.

Becker, B., and M. A. Cooper. 2013. Aminoglycoside antibiotics in the 21st century. ACS chemical biology 8:105–115.

Benveniste, R., and J. Davies. 1973. Structure-activity relationships among the aminoglycoside antibiotics: role of hydroxyl and amino groups. Antimicrobial agents and chemotherapy 4:402–409.

Borovinskaya, M. A., R. D. Pai, W. Zhang, B. S. Schuwirth, J. M. Holton, G. Hirokawa, H. Kaji, A. Kaji, and J. H. D. Cate. 2007. Structural basis for aminoglycoside inhibition of bacterial ribosome recycling. Nature structural & molecular biology 14:727–732.

Böttger, E. C., B. Springer, T. Prammananan, Y. Kidan, and P. Sander. 2001. Structural basis for selectivity and toxicity of ribosomal antibiotics. EMBO reports 2:318–323.

Carter, A. P., W. M. Clemons, D. E. Brodersen, R. J. Morgan-Warren, B. T. Wimberly, and V. Ramakrishnan. 2000. Functional insights from the structure of the 30S ribosomal subunit and its interactions with antibiotics. Nature 407:340–348.

Cortese, J. D., A. L. Voglino, and C. R. Hackenbrock. 1991. Ionic strength of the intermembrane space of intact mitochondria as estimated with fluorescein-BSA delivered by low pH fusion. The Journal of Cell Biology 113:1331–1340.

Dey, D., J. M. Mattingly, N. Zelinskaya, C. M. Dunham, and G. L. Conn. 2025. Basis for selective drug evasion of an aminoglycoside-resistance ribosomal RNA modification. Nature Communications 16:7992.

Ellis, R. J. 2001. Macromolecular crowding: obvious but underappreciated. Trends in Biochemical Sciences 26:597–604.

Emsley, P., B. Lohkamp, W. G. Scott, and K. Cowtan. 2010. Features and development of coot. Acta crystallographica. Section D, Biological crystallography 66:486–501.

Forsberg, B. O., P. N. M. Shah, and A. Burt. 2023. A robust normalized local filter to estimate compositional heterogeneity directly from cryo-EM maps. Nature communications 14:5802.

Fosso, M. Y., Y. Li, and S. Garneau-Tsodikova. 2014. New trends in aminoglycosides use. MedChemComm 5:1075–1091.

Fourmy, D., M. I. Recht, S. C. Blanchard, and J. D. Puglisi. 1996. Structure of the A site of Escherichia coli 16S ribosomal RNA complexed with an aminoglycoside antibiotic. Science (New York, N.Y.) 274:1367–1371.

Ghosh Dastidar, N., N. S. Freyer, V. Petrychenko, A. C. de A P Schwarzer, B.-Z. Peng, E. Samatova, C. Kothe, M. Schmidt, F. Peske, A. Z. Politi, H. Urlaub, N. Fischer, M. V. Rodnina, and I. Wohlgemuth. 2025. Selective silencing of antibiotic-tethered ribosomes as a resistance mechanism against aminoglycosides. Nature Communications 16:9568.

Goddard, T. D., C. C. Huang, E. C. Meng, E. F. Pettersen, G. S. Couch, J. H. Morris, and T. E. Ferrin. 2018. UCSF ChimeraX: Meeting modern challenges in visualization and analysis. Protein science: a publication of the Protein Society 27:14–25.

Greber, B. J., P. Bieri, M. Leibundgut, A. Leitner, R. Aebersold, D. Boehringer, and N. Ban. 2015. Ribosome. The complete structure of the 55S mammalian mitochondrial ribosome. Science (New York, N.Y.) 348:303–308.

Guan, M.-X. 2011. Mitochondrial 12S rRNA mutations associated with aminoglycoside ototoxicity. Mitochondrion 11:237–245.

Hobbie, S. N., S. Akshay, S. K. Kalapala, C. M. Bruell, D. Shcherbakov, and E. C. Böttger. 2008. Genetic analysis of interactions with eukaryotic rRNA identify the mitoribosome as target in aminoglycoside ototoxicity. Proceedings of the National Academy of Sciences of the United States of America 105:20888–20893.

Holm, M., S. K. Natchiar, E. J. Rundlet, A. G. Myasnikov, Z. L. Watson, R. B. Altman, H.-Y. Wang, J. Taunton, and S. C. Blanchard. 2023. mRNA decoding in human is kinetically and structurally distinct from bacteria. Nature 617:200–207.

Holvec, S., C. Barchet, A. Lechner, L. Fréchin, S. N. T. De Silva, I. Hazemann, P. Wolff, O. von Loeffelholz, and B. P. Klaholz. 2024. The structure of the human 80S ribosome at 1.9 Å resolution reveals the molecular role of chemical modifications and ions in RNA. Nature structural & molecular biology 31:1251–1264.

Huang, D. W., B. T. Sherman, and R. A. Lempicki. 2009. Systematic and integrative analysis of large gene lists using DAVID bioinformatics resources. Nature Protocols 4:44–57.

Huang, S., N. A. Aleksashin, A. B. Loveland, D. Klepacki, K. Reier, A. Kefi, T. Szal, J. Remme, L. Jaeger, N. Vázquez-Laslop, A. A. Korostelev, and A. S. Mankin. 2020. Ribosome engineering reveals the importance of 5S rRNA autonomy for ribosome assembly. Nature communications 11:2900.

Keeling, K. M., X. Xue, G. Gunn, and D. M. Bedwell. 2014. Therapeutics based on stop codon readthrough. Annual review of genomics and human genetics 15:371–394.

Kimanius, D., L. Dong, G. Sharov, T. Nakane, and S. H. W. Scheres. 2021. New tools for automated cryo-EM single-particle analysis in RELION-4.0. The Biochemical journal 478:4169–4185.

Kohanski, M. A., D. J. Dwyer, and J. J. Collins. 2010. How antibiotics kill bacteria: from targets to networks. Nature reviews. Microbiology 8:423–435.

Kolosova, O., Y. Zgadzay, A. Stetsenko, A. P. Sukhinina, A. Atamas, S. Validov, A. Rogachev, K. Usachev, L. Jenner, S. E. Dmitriev, G. Yusupova, A. Guskov, and M. Yusupov. 2025. Mechanism of read-through enhancement by aminoglycosides and mefloquine. Proceedings of the National Academy of Sciences of the United States of America 122:e2420261122.

Li, W., S. T.-L. Chang, F. R. Ward, and J. H. D. Cate. 2020. Selective inhibition of human translation termination by a drug-like compound. Nature communications 11:4941.

Li, W., F. R. Ward, K. F. McClure, S. T.-L. Chang, E. Montabana, S. Liras, R. G. Dullea, and J. H. D. Cate. 2019. Structural basis for selective stalling of human ribosome nascent chain complexes by a drug-like molecule. Nature structural & molecular biology 26:501–509.

Li, X., M. Wang, T. Denk, R. Buschauer, Y. Li, R. Beckmann, and J. Cheng. 2024. Structural basis for differential inhibition of eukaryotic ribosomes by tigecycline. Nature communications 15:5481.

Liebschner, D., P. V. Afonine, M. L. Baker, G. Bunkóczi, V. B. Chen, T. I. Croll, B. Hintze, L. W. Hung, S. Jain, A. J. McCoy, N. W. Moriarty, R. D. Oeffner, B. K. Poon, M. G. Prisant, R. J. Read, J. S. Richardson, D. C. Richardson, M. D. Sammito, O. V. Sobolev, D. H. Stockwell, T. C. Terwilliger, A. G. Urzhumtsev, L. L. Videau, C. J. Williams, and P. D. Adams. 2019. Macromolecular structure determination using X-rays, neutrons and electrons: recent developments in Phenix. Acta crystallographica. Section D, Structural biology 75:861–877.

Lin, J., D. Zhou, T. A. Steitz, Y. S. Polikanov, and M. G. Gagnon. 2018. Ribosome-targeting antibiotics: Modes of action, mechanisms of resistance, and implications for drug design. Annual review of biochemistry 87:451–478.

Lintner, N. G., K. F. McClure, D. Petersen, A. T. Londregan, D. W. Piotrowski, L. Wei, J. Xiao, M. Bolt, P. M. Loria, B. Maguire, K. F. Geoghegan, A. Huang, T. Rolph, S. Liras, J. A. Doudna, R. G. Dullea, and J. H. D. Cate. 2017. Selective stalling of human translation through small-molecule engagement of the ribosome nascent chain. PLoS biology 15:e2001882.

Manuvakhova, M., K. Keeling, and D. M. Bedwell. 2000. Aminoglycoside antibiotics mediate context-dependent suppression of termination codons in a mammalian translation system. RNA (New York, N.Y.) 6:1044–1055.

Marín, I., J. P. Abad, D. Ureña, and R. Amils. 1995. High-ionic strength interference of ribosomal inhibition produced by aminoglycoside antibiotics. Biochemistry 34:16519–16523.

Matt, T., C. L. Ng, K. Lang, S.-H. Sha, R. Akbergenov, D. Shcherbakov, M. Meyer, S. Duscha, J. Xie, S. R. Dubbaka, D. Perez-Fernandez, A. Vasella, V. Ramakrishnan, J. Schacht, and E. C. Böttger. 2012. Dissociation of antibacterial activity and aminoglycoside ototoxicity in the 4-monosubstituted 2-deoxystreptamine apramycin. Proceedings of the National Academy of Sciences of the United States of America 109:10984–10989.

Moazed, D., and H. F. Noller. 1987. Interaction of antibiotics with functional sites in 16S ribosomal RNA. Nature 327:389–394.

Ogle, J. M., D. E. Brodersen, W. M. Clemons Jr, M. J. Tarry, A. P. Carter, and V. Ramakrishnan. 2001. Recognition of cognate transfer RNA by the 30S ribosomal subunit. Science (New York, N.Y.) 292:897–902.

Palmer, E., J. M. Wilhelm, and F. Sherman. 1979. Phenotypic suppression of nonsense mutants in yeast by aminoglycoside antibiotics. Nature 277:148–150.

Paternoga, H., C. Crowe-McAuliffe, L. V. Bock, T. O. Koller, M. Morici, B. Beckert, A. G. Myasnikov, H. Grubmüller, J. Nováček, and D. N. Wilson. 2023. Structural conservation of antibiotic interaction with ribosomes. Nature structural & molecular biology 30:1380–1392.

Prezant, T. R., J. V. Agapian, M. C. Bohlman, X. Bu, S. Oztas, W. Q. Qiu, K. S. Arnos, G. A. Cortopassi, L. Jaber, and J. I. Rotter. 1993. Mitochondrial ribosomal RNA mutation associated with both antibiotic-induced and non-syndromic deafness. Nature genetics 4:289–294.

Prokhorova, I., R. B. Altman, M. Djumagulov, J. P. Shrestha, A. Urzhumtsev, A. Ferguson, C.-W. T. Chang, M. Yusupov, S. C. Blanchard, and G. Yusupova. 2017. Aminoglycoside interactions and impacts on the eukaryotic ribosome. Proceedings of the National Academy of Sciences of the United States of America 114:E10899–E10908.

Punjani, A., J. L. Rubinstein, D. J. Fleet, and M. A. Brubaker. 2017. cryoSPARC: algorithms for rapid unsupervised cryo-EM structure determination. Nature methods 14:290–296.

Recht, M. I., S. Douthwaite, and J. D. Puglisi. 1999. Basis for prokaryotic specificity of action of aminoglycoside antibiotics. The EMBO journal 18:3133–3138.

Sabbavarapu, N. M., M. Shavit, Y. Degani, B. Smolkin, V. Belakhov, and T. Baasov. 2016. Design of novel aminoglycoside derivatives with enhanced suppression of diseases-causing nonsense mutations. ACS medicinal chemistry letters 7:418–423.

Shalev, M., H. Rozenberg, B. Smolkin, A. Nasereddin, D. Kopelyanskiy, V. Belakhov, T. Schrepfer, J. Schacht, C. L. Jaffe, N. Adir, and T. Baasov. 2015. Structural basis for selective targeting of leishmanial ribosomes: aminoglycoside derivatives as promising therapeutics. Nucleic acids research 43:8601–8613.

Sherman, B. T., M. Hao, J. Qiu, X. Jiao, M. W. Baseler, H. C. Lane, T. Imamichi, and W. Chang. 2022. DAVID: a web server for functional enrichment analysis and functional annotation of gene lists (2021 update). Nucleic Acids Research 50:W216–W221.

Singh, V., Y. Itoh, S. Del’Olio, A. Hassan, A. Naschberger, R. K. Flygaard, Y. Nobe, K. Izumikawa, S. Aibara, J. Andréll, P. C. Whitford, A. Barrientos, M. Taoka, and A. Amunts. 2024. Mitoribosome structure with cofactors and modifications reveals mechanism of ligand binding and interactions with L1 stalk. Nature communications 15:4272.

Spitzer, J., and B. Poolman. 2009. The role of biomacromolecular crowding, ionic strength, and physicochemical gradients in the complexities of life’s emergence. Microbiology and Molecular Biology Reviews 73:371–388.

Su, B., K. Huang, Z. Peng, A. Amunts, and J. Yang. 2025. CryoAtom improves model building for cryo-EM. Nature structural & molecular biology.

Thompson, R. F., M. G. Iadanza, E. L. Hesketh, S. Rawson, and N. A. Ranson. 2019. Collection, pre-processing and on-the-fly analysis of data for high-resolution, single-particle cryo-electron microscopy. Nature protocols 14:100–118.

Wallis, M. G., U. von Ahsen, R. Schroeder, and M. Famulok. 1995. A novel RNA motif for neomycin recognition. Chemistry & biology 2:543–552.

Wang, L., A. Pulk, M. R. Wasserman, M. B. Feldman, R. B. Altman, J. H. D. Cate, and S. C. Blanchard. 2012. Allosteric control of the ribosome by small-molecule antibiotics. Nature structural & molecular biology 19:957–963.

Wangen, J. R., and R. Green. 2020. Stop codon context influences genome-wide stimulation of termination codon readthrough by aminoglycosides. eLife 9.

Wasserman, M. R., A. Pulk, Z. Zhou, R. B. Altman, J. C. Zinder, K. D. Green, S. Garneau-Tsodikova, J. H. D. Cate, and S. C. Blanchard. 2015. Chemically related 4,5-linked aminoglycoside antibiotics drive subunit rotation in opposite directions. Nature communications 6:7896.

Wilson, D. N. 2014. Ribosome-targeting antibiotics and mechanisms of bacterial resistance. Nature reviews. Microbiology 12:35–48.

Zaher, H. S., and R. Green. 2009. Fidelity at the molecular level: lessons from protein synthesis. Cell 136:746–762.

