## SupplementaryData for "Human ribosomes interactions reveal the binding landscape of neomycin"

[3] Department of Medicinal Chemistry, State Key Laboratory of Discovery and Utilization of Functional Components in Traditional Chinese Medicine, Shandong Key Laboratory of Druggability Optimization and Evaluation for Lead Compounds, Shandong Basic Science Academic Special Zone/Research Center (Pharmacy), School of Pharmaceutical Sciences, Cheeloo College of Medicine, Shandong University, Jinan, Shandong, China

[4] Department of Medical Dataology, School of Public Health, Cheeloo College of Medicine, Shandong University, Jinan, China

[†] Equal contribution: Meng Wu, Lingfan Zhu, Keqin Wu, Changfeng Jia, Xinyi Zhang and Zhongcheng Zhao contributed equally to this work.

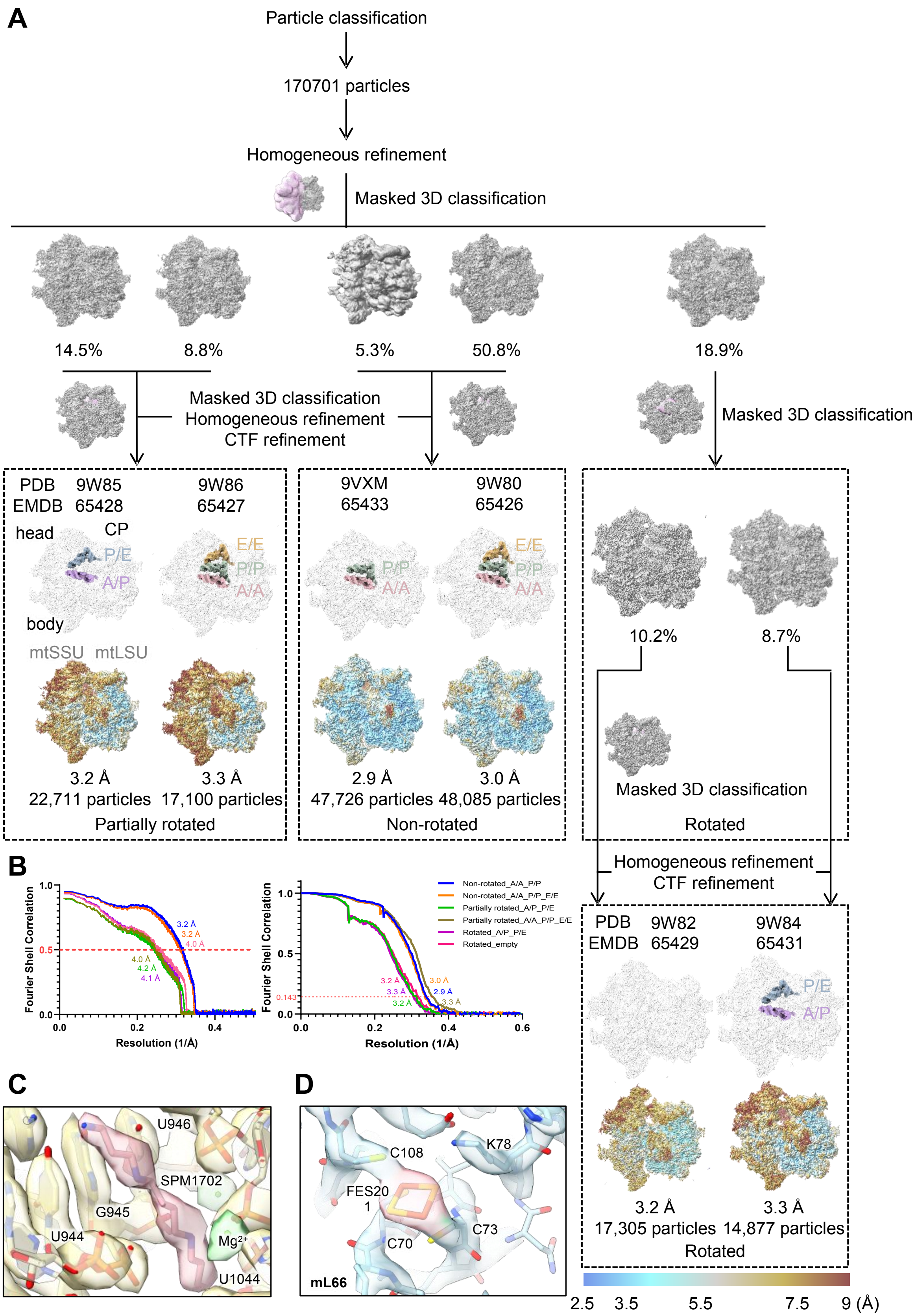
A
Particle classification
170701 particles
Homogeneous refinement
Masked 3D classification
14.5%
8.8%
5.3%
50.8%
18.9%
Masked 3D classification
Homogeneous refinement
CTF refinement
Masked 3D classification
PDB 9W85
EMDB 65428
9W86
65427
head
CP
P/E
A/P
E/E
P/P
A/A
body
mtSSU mtLSU
3.2 Å
3.3 Å
22,711 particles 17,100 particles
Partially rotated
9VXM
65433
9W80
65426
2.9 Å
3.0 Å
47,726 particles 48,085 particles
Non-rotated
10.2%
8.7%
Masked 3D classification
Rotated
Homogeneous refinement
CTF refinement
PDB 9W82
EMDB 65429
9W84
65431
3.2 Å
3.3 Å
17,305 particles 14,877 particles
Rotated
2.5 3.5 5.5 7.5 9 (Å)
B
Fourier Shell Correlation
Resolution (1/Å)
Non-rotated_A/A_P/P
Non-rotated_A/A_P/P_E/E
Partially rotated_A/P_P/E
Partially rotated_A/A_P/P_E/E
Rotated_A/P_P/E
Rotated_empty
0.143
C
U946
SPM1702
G945
U944
Mg2+
U1044
D
C108
K78
FES20
1
C70
C73
mL66

**Figure S1. Cryo-EM data processing of neomycin-treated mitoribosomes**. **(A)** Cryo-EM data processing workflow of mitoribosomes incubated with 50 μM neomycin. Overall structures of mitoribosomes with bound tRNAs and cofactors, along with local resolution estimation results with the resolution scale (Å) are shown. **(B)** Fourier shell correlation (FSC) curves, including map-to-model FSC curves evaluated at the standard FSC = 0.5 threshold (left) and curves for the reported maps indicating overall resolution at FSC = 0.143 (right). **(C-D)** Cryo-EM density for representative cofactors from the same map as Fig. 1 are displayed: SPM1702 **(C)** and FES201 **(D)** bound to the non-rotated mitoribosome with A/A and P/P site tRNA (overall resolution=2.9 Å, σ=0.90).

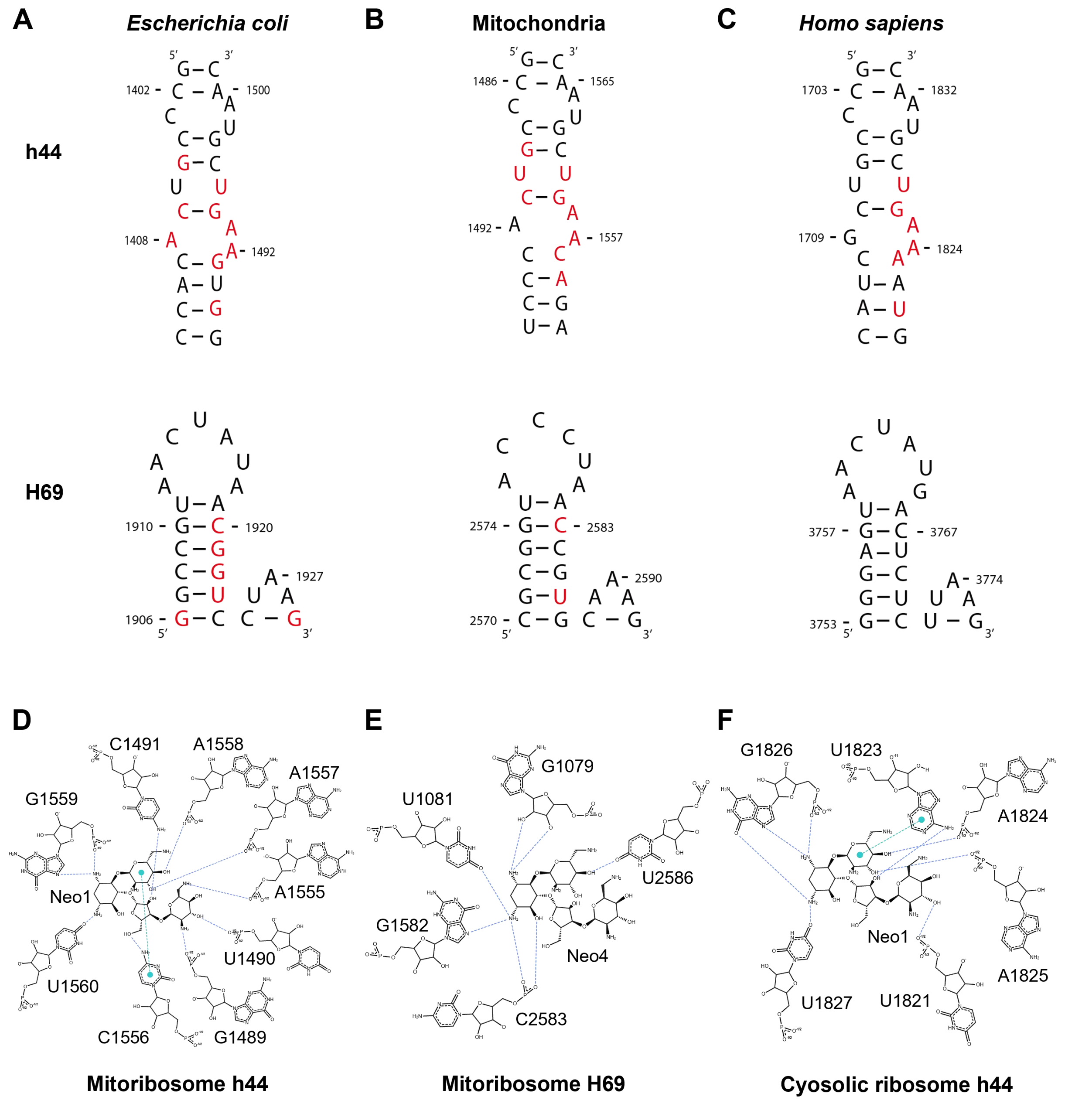


**Figure S2. Comparative analysis of h44 and H69, and neomycin binding. (A-C)** Secondary structures of h44 and H69 from *Escherichia coli* **(A)**, mitochondria **(B),** and *Homo sapiens* cytosolic **(C)** ribosomes. **(D-F)** Schematic representation of h44 and H69 interactions with neomycin.

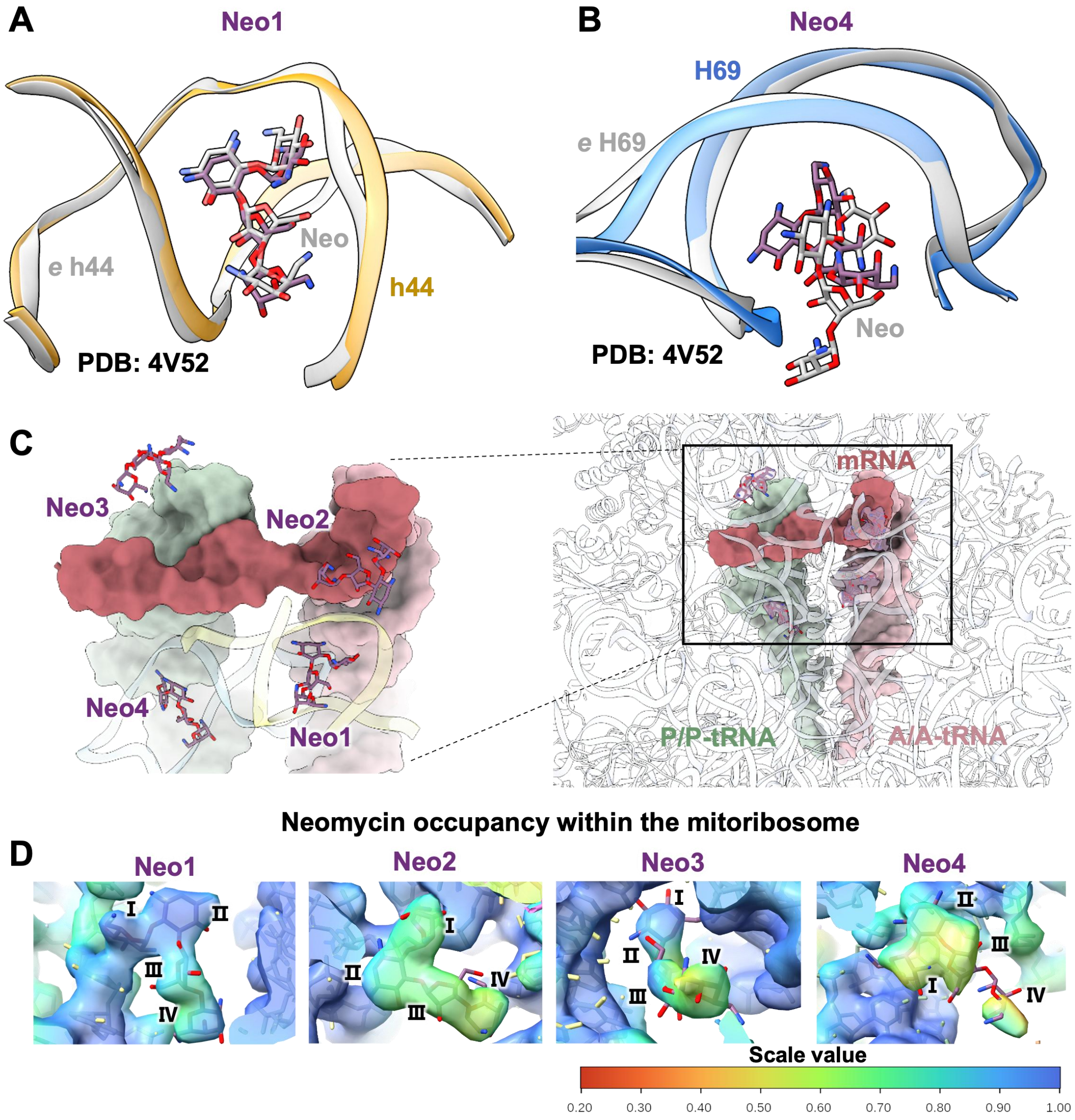


**Figure S3. Analysis of neomycin's binding to mitoribosome's decoding center. (A-B)** Superimposition of neomycin binding pockets within h44 **(A)** and H69 **(B)** with the bacterial ribosome (PDB: 4V52[27]). **(C)** Overview of the mRNA decoding center with bound neomycin molecules. A zoomed inset highlights the bound neomycin molecules. **(D)** Occupancy estimation of Neo1-4 within the decoding center of the mitoribosome. Amplified maps, annotated by the occupancy of neomycin with surrounding ribosomal residues, are colored according to the estimated local scale. The kernel size and radius used were 5 and 2.5 pixels, respectively.

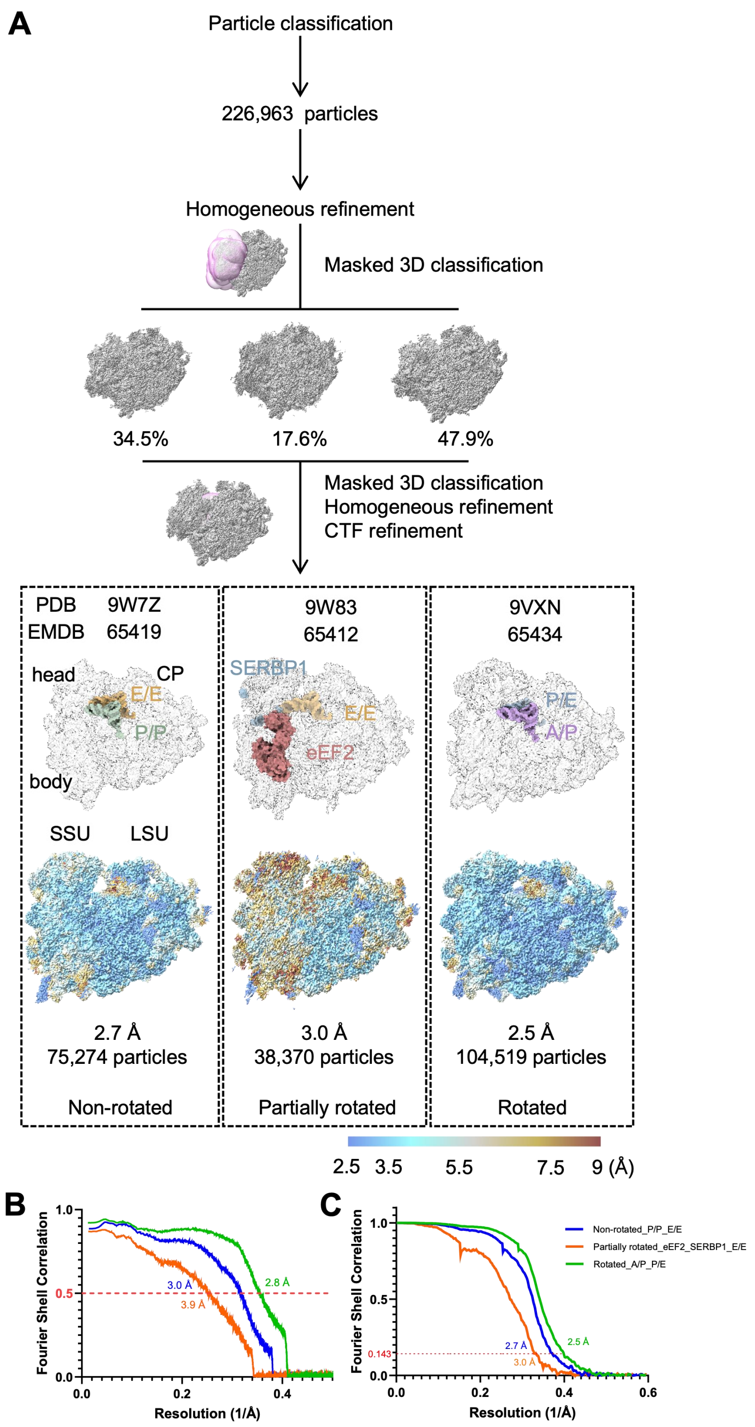


**Figure S4. Cryo-EM data processing of neomycin-treated cytosolic ribosomes**. **(A)** Cryo-EM data processing workflow of cytosolic ribosomes incubated with 50 μM neomycin. Overall structures of cytosolic ribosomes with different bound tRNAs and cofactors, along with local resolution estimation results with the resolution scale in Å are shown. **(B)** FSC curves calculated for the reported maps, indicating an overall resolution at FSC = 0.143. (**C**) FSC curves evaluated between the models and corresponding maps, with the standard FSC = 0.5 threshold indicated.

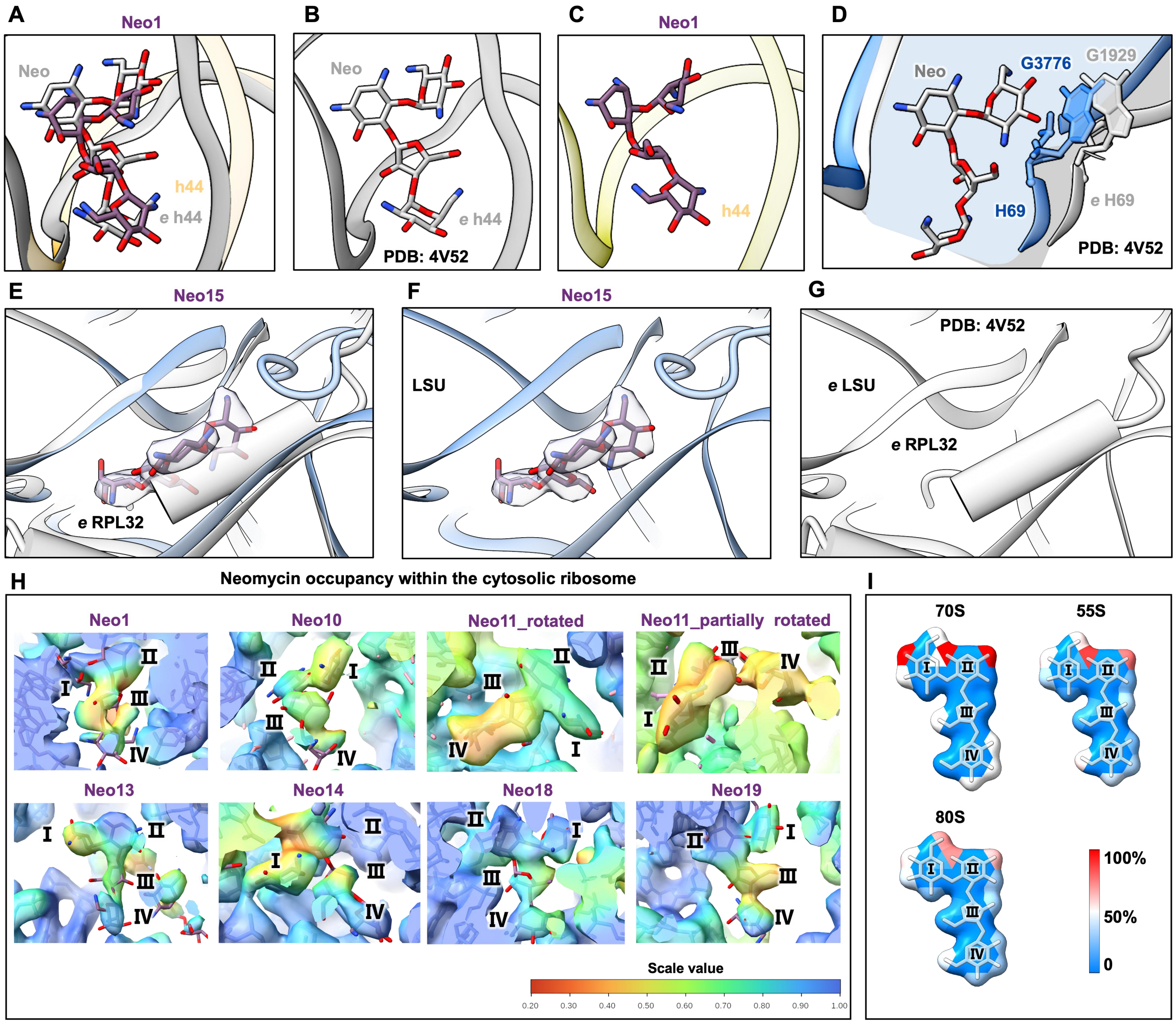


**Figure S5. Structural analysis and occupancy estimation of neomycin's binding sites on the cytosolic ribosome. (A)** Superimposition of the neomycin-bound h44 from this study (yellow) and bacterial ribosome (PDB: 4V52[27], grey). **(B)** Neomycin molecules bound to the bacterial ribosome**. (C)** Neomycin molecules (Neo1) bound to the human cytosolic ribosome. **(D)** Superimposition of H69 from this study (blue) and neomycin-bound H69 (grey) from the bacterial ribosome (PDB: 4V52[27]). Potential steric clashes are highlighted, with contributing nucleotides shown as sticks. The area size of the H69 pocket is indicated in blue (human) and grey (bacteria). **(E)** Superimposition of the neomycin-bound pocket from this study (blue) and bacterial ribosome (PDB: 4V52[27], grey). This neomycin binding pocket is typically occupied by RPL32 protein in the bacterial ribosome. **(F)** Neo15 is shown interacting with this novel cytosolic ribosomal pocket. **(G)** The corresponding pocket is occupied by RPL32 in the bacterial ribosome. **(H)** Occupancy estimation of neomycin molecules within the cytosolic ribosome discussed in the main text. Amplified maps, annotated by the occupancy of neomycin molecules with surrounding ribosomal residues, are colored according to the estimated local scale. The kernel size and radius used were 5 and 2.5 pixels, respectively. **(I)** Conserved interaction modules of neomycin-ribosome complexes. Heatmaps showing the interacting frequency of neomycin rings (I–IV) across bacterial (70S), human mitochondrial (55S), and human cytosolic (80S) ribosomes. The color scale indicates the normalized interaction frequency (0–100%).

77

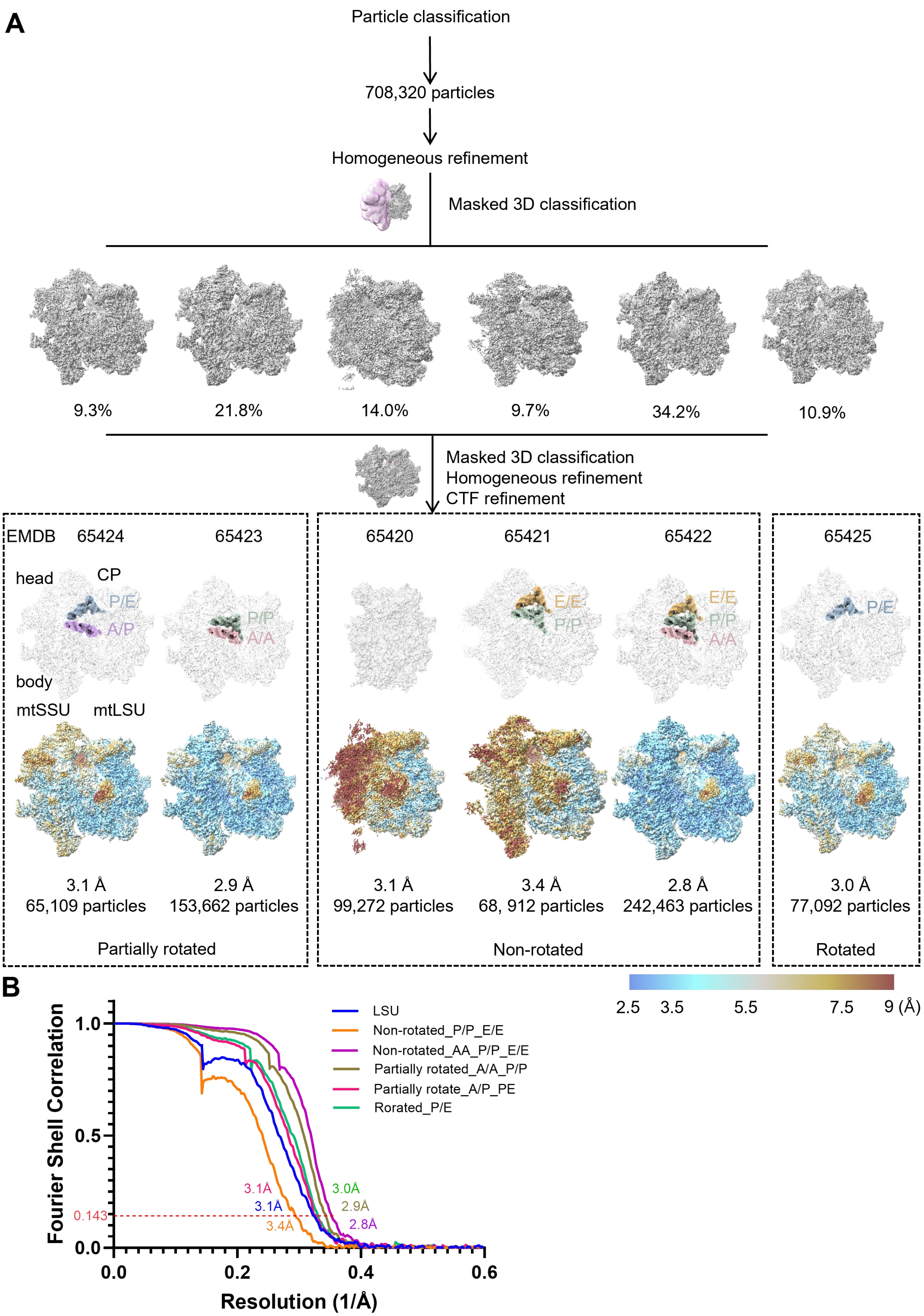
A
Particle classification
708,320 particles
Homogeneous refinement
Masked 3D classification
9.3%
21.8%
14.0%
9.7%
34.2%
10.9%
Masked 3D classification
Homogeneous refinement
CTF refinement
EMDB
65424
65423
65420
65421
65422
65425
head
CP
P/E
A/P
P/P
A/A
E/E
body
mtSSU
mtLSU
3.1 Å
65,109 particles
2.9 Å
153,662 particles
3.1 Å
99,272 particles
3.4 Å
68, 912 particles
2.8 Å
242,463 particles
3.0 Å
77,092 particles
Partially rotated
Non-rotated
Rotated
2.5
3.5
5.5
7.5
9 (Å)
B
Fourier Shell Correlation
Resolution (1/Å)
LSU
Non-rotated_P/P_E/E
Non-rotated_AA_P/P_E/E
Partially rotated_A/A_P/P
Partially rotate_A/P_PE
Rorated_P/E
0.143
3.1Å
3.1Å
3.4Å
3.0Å
2.9Å
2.8Å
1.0
0.5
0.0
0.0
0.2
0.4
0.6

78

**Figure S6. Cryo-EM data processing of mitoribosomes from neomycin-treated cells. (A)** Cryo-EM data processing workflow for native mitoribosomes isolated from 1 mM neomycin-treated cells. Overall structures of mitoribosomes with bound tRNAs and cofactors, along with local resolution estimation results with resolution scale in Å are shown. **(B)** FSC curves calculated for the reported maps, indicating an overall resolution at FSC = 0.143.

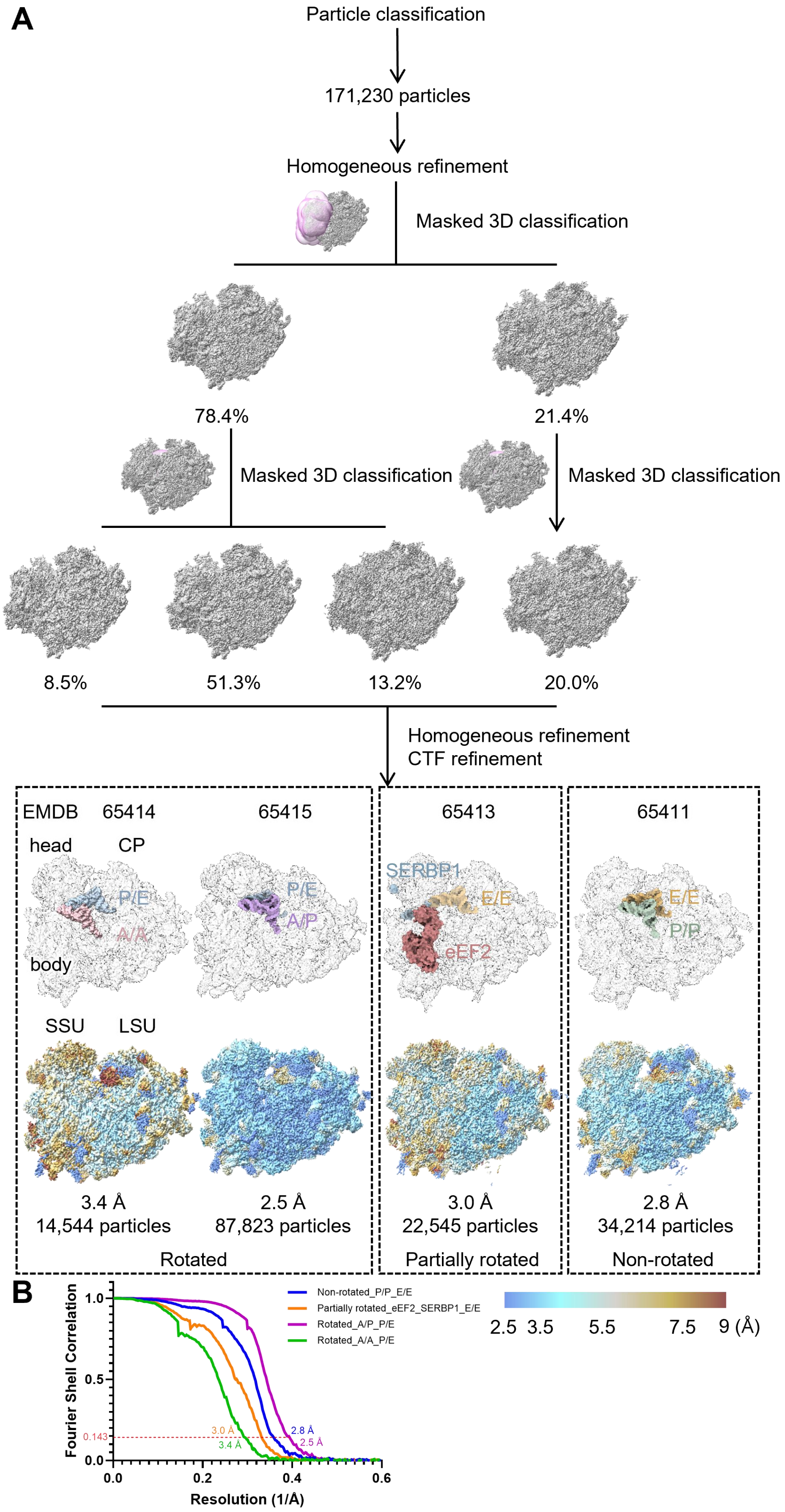
A
Particle classification
171,230 particles
Homogeneous refinement
Masked 3D classification
78.4%
21.4%
Masked 3D classification
Masked 3D classification
8.5%
51.3%
13.2%
20.0%
Homogeneous refinement
CTF refinement
EMDB 65414
65415
65413
65411
head
CP
P/E
A/A
body
P/E
A/P
SERBP1
E/E
eEF2
E/E
P/P
SSU
LSU
3.4 Å
14,544 particles
2.5 Å
87,823 particles
3.0 Å
22,545 particles
2.8 Å
34,214 particles
Rotated
Partially rotated
Non-rotated
B
Fourier Shell Correlation
Resolution (1/Å)
0.143
3.0 Å
3.4 Å
2.8 Å
2.5 Å
Non-rotated_P/P_E/E
Partially rotated_eEF2_SERBP1_E/E
Rotated_A/P_P/E
Rotated_A/A_P/E
2.5 3.5 5.5 7.5 9 (Å)

**Figure S7. Cryo-EM data processing of cytosolic ribosomes from neomycin-treated cells**. **(A)** Cryo-EM data processing workflow for native cytosolic ribosomes isolated from 1 mM neomycin-treated cells. Overall structures of cytosolic ribosomes with different bound tRNAs and cofactors, along with local resolution estimation results with resolution scale in Å are shown. **(B)** FSC curves calculated for the reported maps, indicating an overall resolution at FSC = 0.143.

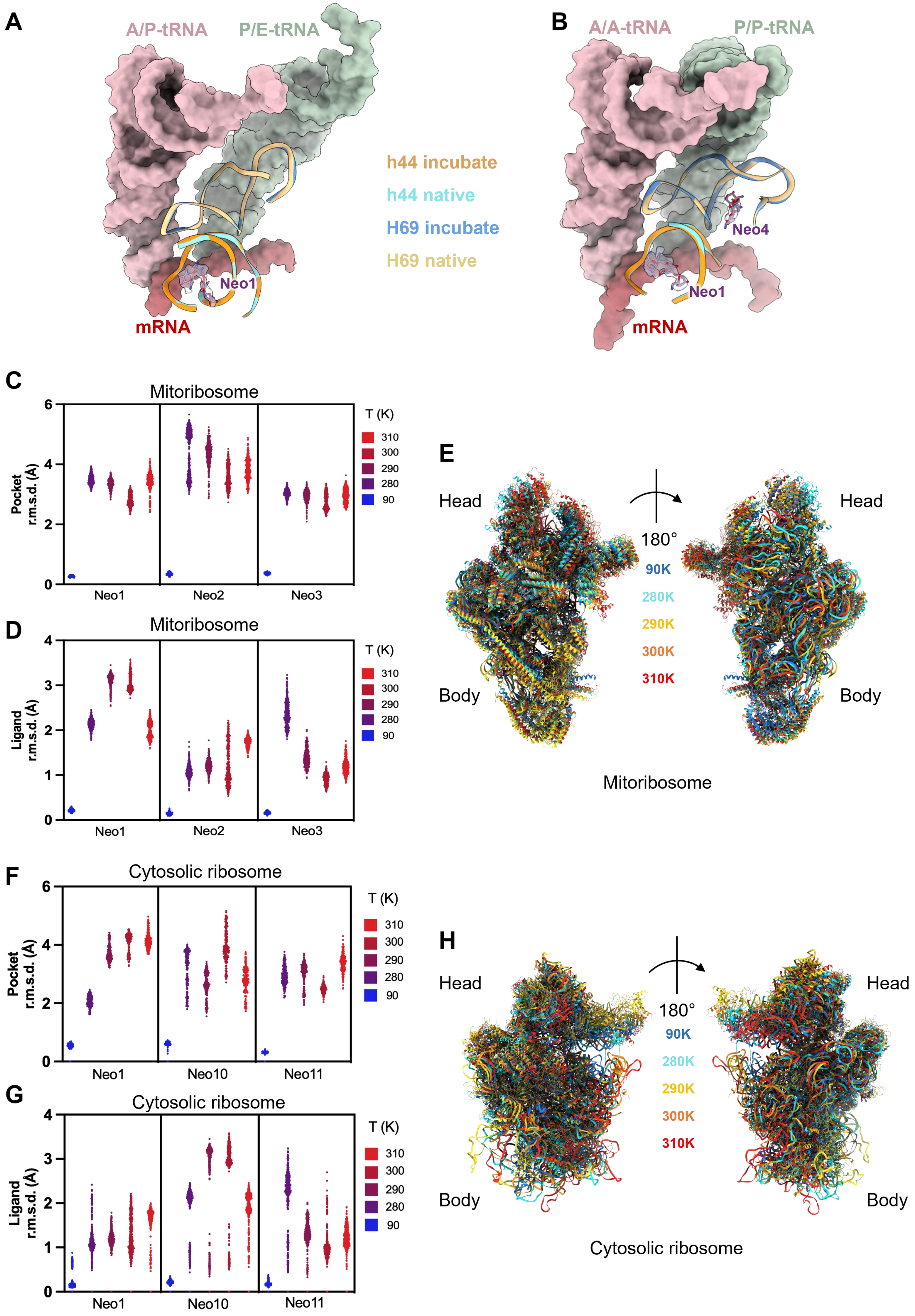
A
A/P-tRNA
P/E-tRNA
h44 incubate
h44 native
H69 incubate
H69 native
Neo1
mRNA
B
A/A-tRNA
P/P-tRNA
Neo4
Neo1
mRNA
C
Mitoribosome
Pocket r.m.s.d. (Å)
Neo1
Neo2
Neo3
T (K)
310
300
290
280
90
D
Mitoribosome
Ligand r.m.s.d. (Å)
Neo1
Neo2
Neo3
E
Head
Body
180°
90K
280K
290K
300K
310K
Mitoribosome
F
Cytosolic ribosome
Pocket r.m.s.d. (Å)
Neo1
Neo10
Neo11
G
Cytosolic ribosome
Ligand r.m.s.d. (Å)
H
Head
Body
Cytosolic ribosome

94

**Figure S8. Structural analysis of the decoding center and MD simulations of neomycin-ribosome interactions and neomycin contact modules. (A-B)** Structure superimposition of decoding centers from the cytosolic ribosome **(A)** and mitoribosome **(B)** following different neomycin treatments. **(C-D)** MD simulations depicting the temperature-dependent binding stability of neomycin molecules (Neo1, Neo2, and Neo3) within mtSSU. The RMSD from their initial binding poses of both the binding pockets **(C)** and the neomycin molecules **(D),** across temperatures from 90 K to 310 K. **(E)** Representative mtSSU structures generated by temperature-dependent MD simulations. **(F-G)** MD simulations depicting the temperature-dependent binding stability of neomycin molecules (Neo1, Neo10, and Neo11) within cytosolic SSU. The RMSD from their initial binding poses of both binding pockets **(F)** and their initial binding poses for neomycin molecules **(G)** are shown across temperatures from 90 K to 310 K. **(H)** Representative cytosolic SSU structures generated by temperature-dependent MD simulations.

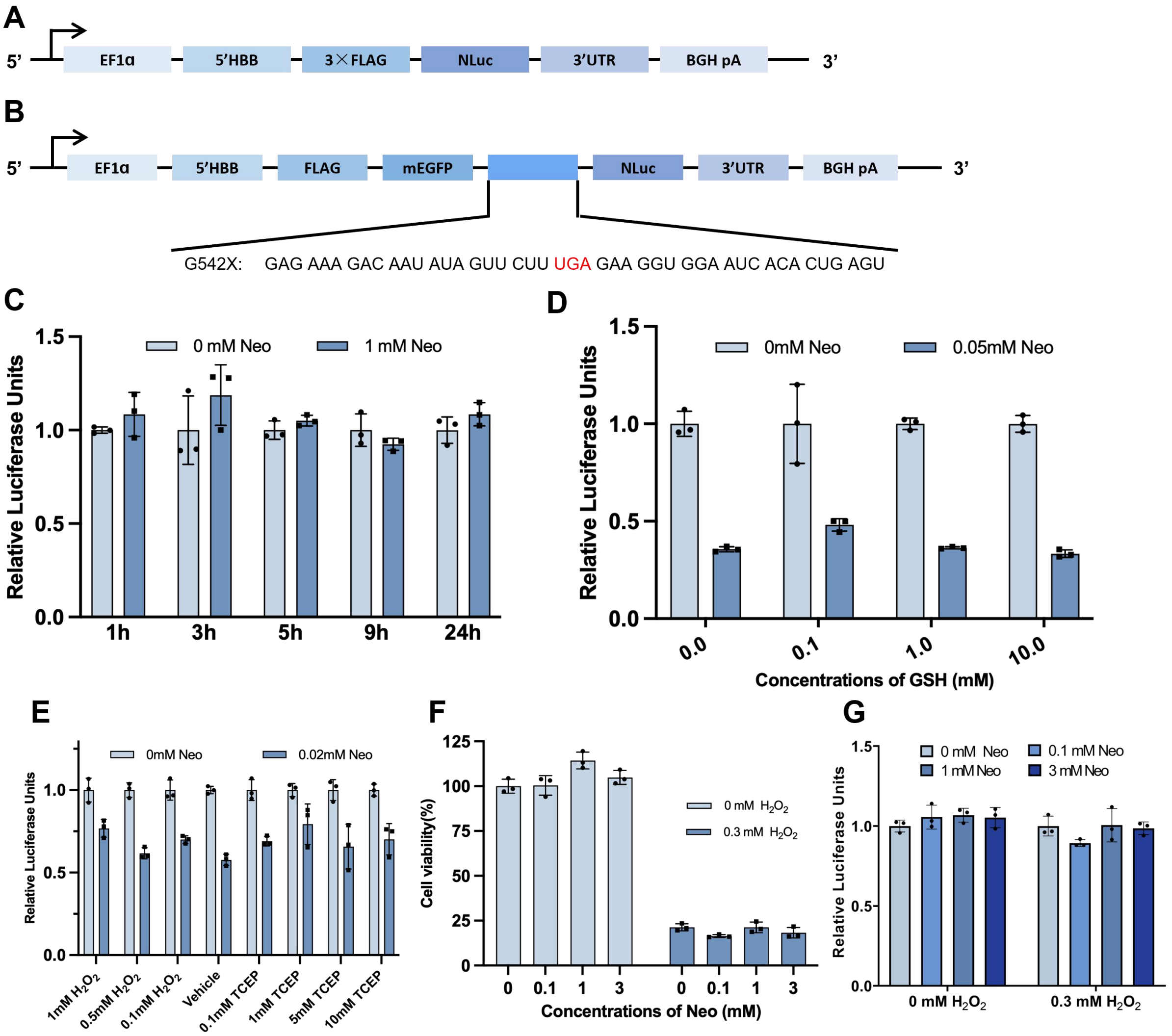


**Figure S9. *In vitro* and *in-cell* translation assays. (A-B)** Schematic representation of the NLuc (**A**) and CFTR (G542X) (**B**) lentiviral constructs used to create stable cell lines. **(C)** Luciferase reporter assays using the NLuc cell line at various time points following treatment with different concentrations of neomycin. **(D)** *In vitro* translation assays with luciferase reporters in response to different concentrations of neomycin and increased concentrations of 0.1, 1 and 10 mM GSH. **(E)** *In vitro* translation assays with luciferase reporters were performed in the presence of 0.02 mM neomycin under varying redox conditions. The redox conditions are mimicked by adding different concentrations of $H_2O_2$ and TCEP. The vehicle group presents the condition with no additional redox agents (data are presented as mean ± s.d., n = 3 independent experiments). **(F)** Cell viability of HEK293T cells measured by the CCK-8 assay after treatment with increasing concentrations of neomycin in the presence or absence of $H_2O_2$ (data are presented as mean ± s.d., n = 3 independent experiments). **(G)** *In-cell* translation assays with luciferase reporters in response to different concentrations of neomycin in a 0.3 mM $H_2O_2$ environment (data are presented as mean ± s.d., n = 3 independent experiments).

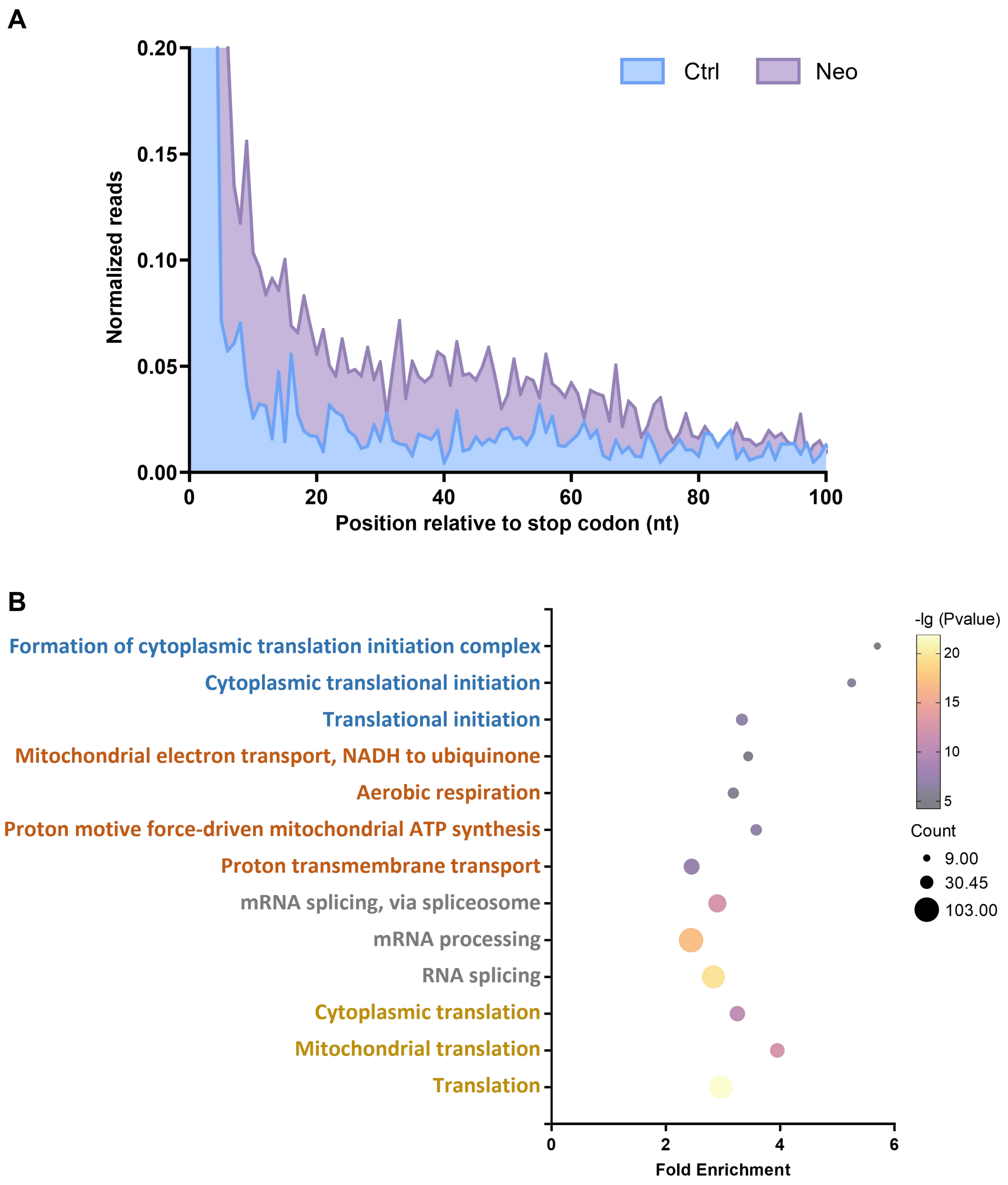


**Figure S10. Ribosome profiling analysis of neomycin-induced translational fidelity defects. (A)** Metagene analysis of ribosome occupancy. Average gene plots showing normalized ribosome footprint densities relative to the distance from the canonical stop codon. Data are compared between untreated (Ctrl) and neomycin-treated (2 mg/mL, ~2.2 mM; 24 h) HEK293T cells. **(B)** Functional clustering of genes exhibiting neomycin-induced readthrough. Cluster analysis performed on 2,102 identified genes, comprising de novo readthrough candidates and those exhibiting at least a two-fold increase in RRTS following neomycin treatment. The plot displays Biological Process (BP) terms for the top four enrichment clusters. Terms within the same functional cluster are indicated by identical colors. Fold Enrichment= (Count/List Total)/(Pop Hits/Pop Total)

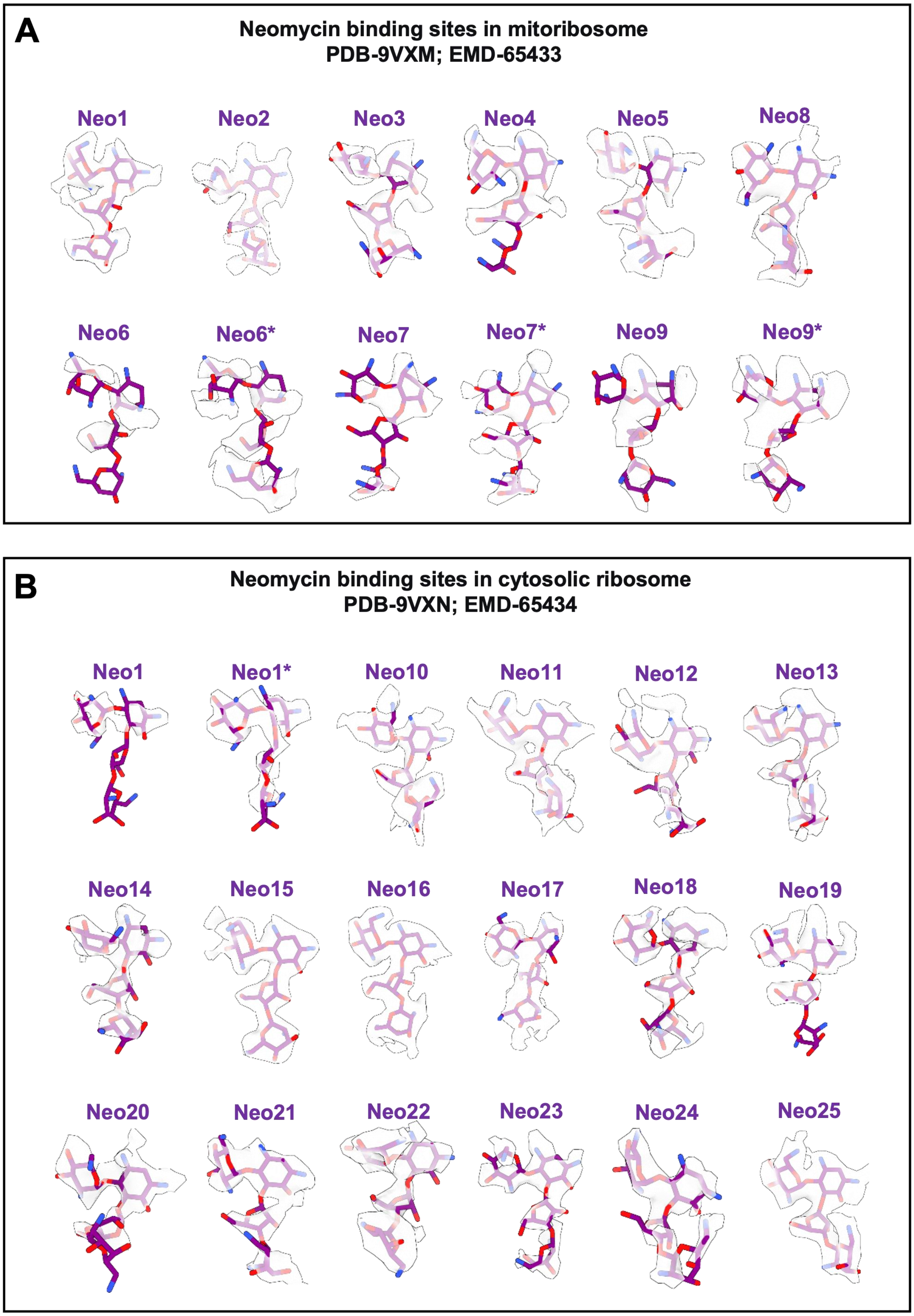
A
Neomycin binding sites in mitoribosome
PDB-9VXM; EMD-65433
Neo1
Neo2
Neo3
Neo4
Neo5
Neo8
Neo6
Neo6*
Neo7
Neo7*
Neo9
Neo9*
B
Neomycin binding sites in cytosolic ribosome
PDB-9VXN; EMD-65434
Neo1
Neo1*
Neo10
Neo11
Neo12
Neo13
Neo14
Neo15
Neo16
Neo17
Neo18
Neo19
Neo20
Neo21
Neo22
Neo23
Neo24
Neo25

**Figure S11. Model and map of Neo bound to ribosomes after *in vitro* incubation with neomycin.**

**(A)** Model and map of Neo bound to the mitoribosome.

**(B)** Model and map of Neo bound to the cytosolic ribosome.

**Table S1. Cryo-EM data collection, processing, model refinement, and validation statistics for mitoribosomes incubated with 50 μM neomycin.**

| | Mitoribosome, Non-rotated state, A/A-P/P | Mitoribosome, Non-rotated state, A/A-P/P-E/E | Mitoribosome, Partially rotated state, A/A-P/P-E/E | Mitoribosome, Partially rotated state, A/P-P/E | Mitoribosome, Rotated state | Mitoribosome, Rotated state, A/P-P/E |
|---|---|---|---|---|---|---|
| EMDB | 65433 | 65426 | 65427 | 65428 | 65429 | 65431 |
| PDB | 9VXM | 9W80 | 9W86 | 9W85 | 9W82 | 9W84 |
| **Data collection and processing** | | | | | | |
| Magnification | 96000x | 96000x | 96000x | 96000x | 96000x | 96000x |
| Voltage (kV) | 300 | 300 | 300 | 300 | 300 | 300 |
| Electron exposure (e–/Å$^2$) | 50 | 50 | 50 | 50 | 50 | 50 |
| Defocus range (μm) | -1 to -2 | -1 to -2 | -1 to -2 | -1 to -2 | -1 to -2 | -1 to -2 |
| Pixel size (Å) | 0.81 | 0.81 | 0.81 | 0.81 | 0.81 | 0.81 |
| Symmetry imposed | *C1* | *C1* | *C1* | *C1* | *C1* | *C1* |
| Initial particle images (no.) | 402,000 | 402,000 | 402,000 | 402,000 | 402,000 | 402,000 |
| Final particle images (no.) | 47,726 | 48,085 | 17,100 | 22,711 | 17,305 | 14,877 |
| Map resolution (Å) | 2.9 | 3.0 | 3.3 | 3.2 | 3.2 | 3.3 |
| FSC threshold | 0.143 | 0.143 | 0.143 | 0.143 | 0.143 | 0.143 |
| Map resolution range (Å) | 2.3-9.2 | 2.4-9.6 | 2.7-11.3 | 2.4-10.7 | 2.8-10.3 | 2.8-10.5 |
| **Refinement** | | | | | | |
| Initial model used (PDB code) | 7QI5 | 7QI5 | 7QI5 | 7QI5 | 7QI5 | 7QI5 |
| Model resolution (Å) | 3.2 | 3.2 | 4.0 | 4.2 | 4.0 | 4.1 |
| FSC threshold | 0.5 | 0.5 | 0.5 | 0.5 | 0.5 | 0.5 |
| Map sharpening *B* factor (Å$^2$) | 58.6 | 56.5 | 29.2 | 31.2 | 31.7 | 33.5 |
| Model composition | | | | | | |
| Non-hydrogen atoms | 176,080 | 177,521 | 177,396 | 175,553 | 172,574 | 175,667 |
| Protein residues | 14,362 | 14,362 | 14,362 | 14,336 | 14,336 | 14,342 |
| Ligands | 285 | 285 | 283 | 277 | 278 | 277 |
| *B* factors (Å$^2$) | | | | | | |
| Protein | 126.12 | 126.12 | 137.36 | 166.45 | 101.27 | 87.98 |
| Ligand | 60.26 | 60.26 | 70.17 | 80.9 | 53.02 | 49.28 |
| R.m.s. deviations | | | | | | |
| Bond lengths (Å) | 0.004 | 0.003 | 0.004 | 0.005 | 0.005 | 0.005 |
| Bond angles (°) | 0.839 | 0.592 | 0.832 | 0.597 | 0.853 | 0.853 |
| Validation | | | | | | |
| MolProbity score | 1.60 | 1.66 | 1.56 | 1.56 | 1.5 | 1.55 |
| Clashscore | 7.51 | 8.46 | 8.1 | 9.24 | 8.14 | 9.83 |
| Poor rotamers (%) | 1.78 | 1.84 | 1.24 | 1.24 | 1.19 | 1.19 |
| Ramachandran plot | | | | | | |
| Favored (%) | 98.36 | 98.32 | 97.79 | 98.01 | 98.44 | 98.44 |
| Allowed (%) | 1.62 | 1.67 | 2.2 | 1.98 | 1.55 | 1.55 |
| Disallowed (%) | 0.01 | 0.01 | 0.01 | 0.01 | 0.01 | 0.01 |

**Table S2**. **Neomycin binding pockets in *in vitro* and *in-cell* mitoribosome states.** This table details the neomycin molecules bound in different mitoribosome states, including their respective binding pockets and proximal nucleotide information. Asterisks (*) denote the quality of the neomycin density, where the number of asterisks corresponds to the count of discernible organic rings of neomycin in the density map.

| State | | | *In vitro* | | | | | | *In-cell* | | | | | |
|---|---|---|---|---|---|---|---|---|---|---|---|---|---|---|
| Number | Binding pocket | Proximal nucleotide | Mitoribosome, Non-rotated state, A/A-P/P | Mitoribosome, Non-rotated state, A/A-P/P-E/E | Mitoribosome, Partially rotated state, A/A-P/P-E/E | Mitoribosome, Partially rotated state, A/P-P/E | Mitoribosome, Rotated state, A/P-P/E | Mitoribosome, Rotated state | mtLSU | Mitoribosome, Non-rotated state, P/P-E/E | Mitoribosome, Non-rotated state, A/A-P/P-E/E | Mitoribosome, Partially rotated state, A/A-P/P | Mitoribosome, Partially rotated state, A/P-P/E | Mitoribosome, Rotated state, P/E |
| Neo1 | h44 | 12S A1556 | **** | **** | *** | *** | ** | ** | | | | | | |
| Neo2 | h1+h44 | 12S C1487 | **** | **** | *** | *** | *** | *** | | | | | | |
| Neo3 | h28 | 12S A1470 | **** | **** | *** | **** | **** | **** | | | | | | |
| Neo4 | H69 | 16S C2583 | *** | *** | | | | | | | | | | |
| Neo5 | H91+H95 | 16S G3019 | **** | **** | **** | | **** | **** | | | | | | |
| Neo6 | H35# | 16S G1949 | ** | ** | *** | | | | | | | | | |
| Neo7 | H33# | 16S U2404 | **** | *** | | ** | | **** | | | | | | |
| Neo8 | H73+H72 | 16S A2707 | **** | **** | *** | **** | **** | **** | | | | | | |
| Neo9 | H39+H37+H75 | 16S C2116 | ** | ** | | ** | ** | ** | | | | | | |

#: rRNA has relatively low conservation.

**Table S3**. **Cryo-EM data collection, processing, model refinement, and validation statistics for cytosolic ribosomes incubated with 50 μM neomycin.**

| | Cytosolic ribosome, Rotated state, A/P-P/E | Cytosolic ribosome, Partially rotated state, eEF2, SERBP1, E/E | Cytosolic ribosome, Non-rotated state, P/P-E/E |
|---|---|---|---|
| EMDB | 65434 | 65412 | 65419 |
| PDB | 9VXN | 9W83 | 9W7Z |
| **Data collection and processing** | | | |
| Magnification | 96000x | 96000x | 96000x |
| Voltage (kV) | 300 | 300 | 300 |
| Electron exposure (e–/Å$^2$) | 50 | 50 | 50 |
| Defocus range (μm) | -1 to -2 | -1 to -2 | -1 to -2 |
| Pixel size (Å) | 0.81 | 0.81 | 0.81 |
| Symmetry imposed | *C1* | *C1* | *C1* |
| Initial particle images (no.) | 972,883 | 972,883 | 972,883 |
| Final particle images (no.) | 104,519 | 38,370 | 75,274 |
| Map resolution (Å) | 2.5 | 3.2 | 2.7 |
| FSC threshold | 0.143 | 0.143 | 0.143 |
| Map resolution range (Å) | 2.0-7.5 | 2.3-9.8 | 2.0-8.6 |
| **Refinement** | | | |
| Initial model used (PDB code) | 8QOI | 8QOI | 8QOI |
| Model resolution (Å) | 2.8 | 3.9 | 3.0 |
| FSC threshold | 0.5 | 0.5 | 0.5 |
| Map sharpening *B*factor (Å$^2$) | 38.8 | 38.4 | 39.6 |
| Model composition | | | |
| Non-hydrogen atoms | 218,051 | 222,621 | 217,694 |
| Protein residues | 11,439 | 12,250 | 11,439 |
| Ligands | 429 | 426 | 426 |
| *B* factors (Å$^2$) | | | |
| Protein | 99.12 | 101.04 | 94.31 |
| Ligand | 42.64 | 47.53 | 43.65 |
| R.m.s. deviations | | | |
| Bond lengths (Å) | 0.004 | 0.062 | 0.006 |
| Bond angles (°) | 0.848 | 1.065 | 0.797 |
| Validation | | | |
| MolProbity score | 1.81 | 2.41 | 2.14 |
| Clashscore | 8.87 | 13.06 | 9.17 |
| Poor rotamers (%) | 1.41 | 3.66 | 3.16 |
| Ramachandran plot | | | |
| Favored (%) | 96.55 | 94.88 | 96.09 |
| Allowed (%) | 3.29 | 4.99 | 3.73 |
| Disallowed (%) | 0.16 | 0.13 | 0.19 |

**Table S4**. **Neomycin binding pockets in cytosolic ribosome states.** This table details the neomycin molecules bound in various cytosolic ribosome states, including their respective binding pockets and proximal nucleotide information. Asterisks (*) denote the quality of the neomycin density, where the number of asterisks corresponds to the count of discernible organic rings of neomycin in the density map.

| State | | | *In vitro* | | | *In-cell* | | | |
|---|---|---|---|---|---|---|---|---|---|
| Number | Binding pocket | Proximal nucleotide | Cytosolic ribosome, Non-rotated state, P/P-E/E | Cytosolic ribosome, Partially rotated state, eEF2, SERBP1, E/E | Cytosolic ribosome, Rotated state, A/P-P/E | Cytosolic ribosome, Non-rotated state, P/P-E/E | Cytosolic ribosome, Partially rotated state, eEF2, SERBP1, E/E | Cytosolic ribosome, Rotated state, A/A-P/E | Cytosolic ribosome, Rotated state, A/P-P/E |
| Neo1 | h44 | 18S A1825 | ** | ** | *** | | | | |
| Neo10 | h28+h37+h40 | 18S U1343 | **** | | **** | **** | **** | | **** |
| Neo11 | h44 | 18S G1726 | **** | **** | **** | | ** | ** | **** |
| Neo12 | H62+H34 | 28S G2868 | **** | **** | **** | **** | | ** | **** |
| Neo13 | H41+H42+H89 | 28S A1939 | **** | **** | **** | **** | | **** | **** |
| Neo14 | H89+H10+H72 | 28S A2047 | **** | *** | **** | | | | *** |
| Neo15 | H24+H26+H2（23S in Prokaryote，28S+5.8S in Eukaryotes） | 28S G2358 | **** | **** | **** | **** | **** | **** | **** |
| Neo16 | H35+H33 | 28S U1616 | **** | **** | **** | **** | **** | **** | **** |
| Neo17 | H39+5s rRNA | 28S U1861 | **** | **** | **** | **** | **** | **** | **** |
| Neo18 | H42 | 28S C1931 | **** | **** | **** | **** | **** | **** | **** |
| Neo19 | H54+H53+L7a | 28S G2777 | **** | ** | **** | | | **** | |
| Neo20 | H58# | 28S G2645 | **** | *** | **** | ** | | | ** |
| Neo21 | H91+H95 | 28S U4481 | **** | **** | **** | | | | |
| Neo22 | H58 | 28S C2683 | **** | **** | **** | | | | **** |
| Neo23 | H74 | 28S C3909 | | ** | **** | | | | ** |
| Neo24 | H89+H91+H42+H90 | 28S A4484 | | | **** | | | | |
| Neo25 | H66+h24（The interface of the LSU and SSU） | 28S C3696 | | **** | **** | | **** | **** | **** |

#: rRNA has relatively low conservation

**Table S5. Cryo-EM data collection and processing statistics for mitoribosomes and cytosolic ribosomes isolated from neomycin-treated cells.**

| **Mitoribosomes from 1mM Neo treated cells** | mtLSU | Mitoribosome, Non-rotated state, P/P-E/E | Mitoribosome, Non-rotated state, A/A-P/P-E/E | Mitoribosome, Partially rotated state, A/A-P/P | Mitoribosome, Partially rotated state, A/P-P/E | Mitoribosome, Rotated state, P/E |
|---|---|---|---|---|---|---|
| EMDB | 65420 | 65421 | 65422 | 65423 | 65424 | 65425 |
| **Data collection and processing** | | | | | | |
| Magnification | 96000x | 96000x | 96000x | 96000x | 96000x | 96000x |
| Voltage (kV) | 300 | 300 | 300 | 300 | 300 | 300 |
| Electron exposure (e–/Å$^2$) | 50 | 50 | 50 | 50 | 50 | 50 |
| Defocus range (μm) | -1 to -2 | -1 to -2 | -1 to -2 | -1 to -2 | -1 to -2 | -1 to -2 |
| Pixel size (Å) | 0.81 | 0.81 | 0.81 | 0.81 | 0.81 | 0.81 |
| Symmetry imposed | *C1* | *C1* | *C1* | *C1* | *C1* | *C1* |
| Initial particle images (no.) | 1,043,603 | 1,043,603 | 1,043,603 | 1,043,603 | 1,043,603 | 1,043,603 |
| Final particle images (no.) | 99,272 | 68,912 | 242,463 | 153,662 | 65,109 | 77,092 |
| Map resolution (Å)<br>FSC threshold | 3.1<br>0.143 | 3.4<br>0.143 | 2.8<br>0.143 | 2.9<br>0.143 | 3.1<br>0.143 | 3.0<br>0.143 |
| Map resolution range (Å) | 2.5-9.2 | 2.5-10.4 | 2.2-6.8 | 2.4-9.2 | 2.6-9.3 | 2.4-10.2 |

| **Cytosolic ribosomes from 1mM Neo treated cells** | Cytosolic ribosome, Non-rotated state, P/P-E/E | Cytosolic ribosome, Partially rotated state, eEF2, SERBP1, E/E | Cytosolic ribosome, Rotated state, A/A-P/E | Cytosolic ribosome, Rotated state, A/P-P/E |
|---|---|---|---|---|
| EMDB | 65411 | 65413 | 65414 | 65415 |
| **Data collection and processing** | | | | |
| Magnification | 96000x | 96000x | 96000x | 96000x |
| Voltage (kV) | 300 | 300 | 300 | 300 |
| Electron exposure (e–/Å$^2$) | 50 | 50 | 50 | 50 |
| Defocus range (μm) | -1 to -2 | -1 to -2 | -1 to -2 | -1 to -2 |
| Pixel size (Å) | 0.81 | 0.81 | 0.81 | 0.81 |
| Symmetry imposed | *C1* | *C1* | *C1* | *C1* |
| Initial particle images (no.) | 338,609 | 338,609 | 338,609 | 338,609 |
| Final particle images (no.) | 34,214 | 22,545 | 14,544 | 87,823 |
| Map resolution (Å)<br>FSC threshold | 2.8<br>0.143 | 3.0<br>0.143 | 3.4<br>0.143 | 2.5<br>0.143 |
| Map resolution range (Å) | 2.3-9.6 | 2.5-9.4 | 2.7-11.3 | 2.3-6.7 |
